# Human hippocampus and dorsomedial prefrontal cortex infer and update latent causes during social interaction

**DOI:** 10.1101/2023.09.21.558855

**Authors:** Ali Mahmoodi, Shuyi Luo, Caroline Harbison, Payam Piray, Matthew Rushworth

## Abstract

When an important event occurs, the observer should identify which features of the environment might have caused it. This is the latent cause inference problem, and it must be solved if observers are to understand their environments. The problem is acute in social settings where individuals may not make equal contributions to the outcomes they achieve together. Here, we designed a novel task in which participants inferred which of two characters was the more likely to have been responsible for outcomes achieved by working together. Using computational modelling, univariate and multivariate analysis of human fMRI, and continuous theta burst stimulation we identified two brain regions that solved the task. Notably, as each outcome occurred, it was possible to decode inference of its cause (the responsible character) from activity in hippocampus. Activity in dorsomedial prefrontal cortex updated estimates of association between cause -- the responsible character – and the outcome.

## Introduction

Latent cause inference is the process by which we infer the causes of our observations^1^. The causes of the events we observe in the world are often ambiguous and may not be directly observable themselves; instead they need to be discovered or inferred^2–5^. Therefore, to learn from our experiences, we first need to identify the causes responsible for those experiences. Identifying the most relevant (causal) determinants of an event is of particular importance in our social life^6^, as most of our observations concern groups of people, rather than single individuals: football players win tournaments within a team in the same way that scientists co-author academic publications together. In these situations, we might not hold all the football players or the scientists equally responsible for the outcomes they achieve together. We might instead identify particular individuals as being especially responsible for the outcomes. Moreover, once we have inferred which individuals are most likely to have been responsible, we update our estimate of their abilities in the light of the new trophy or publication more than we do for those individuals we believe have contributed very little to those outcomes.

In summary, the first problem an observer faces when witnessing an important event is to infer the latent cause in the environment that was responsible for the outcome. This is then followed by a second process: the observer updates their estimate of the strength of the association between the inferred cause and the observed outcome. We refer to the first process as latent cause inference (LCI) and the second process as updating estimates of association (UEA) between cause and outcome. This two-stage computational process might therefore engage not one brain area but two or more areas comprising a neural circuit.

LCI and UEA may depend on different neural mechanisms. Hippocampus is one of the two areas on which we focus. It is established that hippocampus supports context-dependent learning^7,8^, navigation^9^, and configural learning^10^. One interpretation of such findings is that these tasks entail inference and hippocampus is concerned with inferring the latent state^11,122,4,13^. However, direct evidence for the role of hippocampus in inferring latent state is currently missing. We show that LCI in fact depends on hippocampus: hippocampus constructs a continually updating map of a social space defined by the relative abilities of different individuals. Second, we show that this allows hippocampus to perform latent state inference in the social domain. Finally, we capitalize on the brain’s specialization for the perception of social stimuli (the specialization of areas such as the fusiform face area [FFA]) to show that when social identities are perceived they are represented in FFA and dmPFC, but when they are inferred, they are represented in hippocampus. We examine how hippocampus constructs such representations through interactions with dmPFC.

We turned to social situations in which multiple characters interact in different ways on different occasions, as they provide an ideal context for investigating LCI and UEA related computations. Notably, despite the ubiquity of LCI in social situations, the investigation of social cognition has been dominated by studies lacking a requirement for inference, even though learning has been extensively investigated^14^. Learning and representing others’ abilities in these situations are supported by a network of brain areas including dorsomedial prefrontal cortex (dmPFC), temporoparietal junction (TPJ), and precuneus^15–18^. Therefore, UEA related computations might be carried out by one or more of these areas instead of, or in conjunction with, hippocampus.

Here we develop an experimental framework for studying how the human brain performs LCI and UEA in the social environment. In each trial, while undergoing functional magnetic resonance imaging (fMRI), participants observed an outcome that they believed was achieved by a pair of characters working together. The participants’ task was to indicate which character was more responsible for a given outcome. Behavioural data analysis and computational modelling indicated that participants did not infer that each outcome was caused to an equal degree by all the characters (the potential latent causes) present in each trial of the task. Instead, they identified one character as being more responsible. Moreover, once they had identified this character (LCI), they updated their estimate of the association (UEA) between that character and the outcome more than they updated their estimate of the association between the other character and the outcome. In the context of this task, the association between a character and an outcome might intuitively be thought of as the character’s ability; when participants perform UEA in this task they are updating an estimate of the character’s ability. As a consequence of this asymmetric UEA process, we show that participants updated their estimate of the best character more following good outcomes, and they updated their estimate of the worst character more following poor outcomes. At a neural level, we identified activity patterns spanning two brain areas that mediated the two-stage inference and updating process. First, hippocampus identified the latent causes of outcomes (LCI); remarkably decoding of hippocampal activity revealed which cause (which character) each participant inferred as the responsible one in each social situation as soon as the outcome was witnessed. Second dmPFC activity mediated the asymmetric updating of estimates of association (UEA). Consistent with this result, temporary disruption of dmPFC activity using continuous theta burst stimulation (cTBS) impaired UEA; it impaired updating of estimates of characters responsibilities for outcomes on the basis of LCI. However, LCI itself remained intact.

## Results

### Experimental task

In experiment 1, participants (*n*=33) performed a latent social cause inference task and a localiser task while undergoing functional magnetic resonance imaging (fMRI; Figure 1a-b, see Methods for details). Participants were told that three characters were paired together in various groups of two to perform a task for several trials. In each trial, participants observed an outcome that they believed had been achieved by one of the pairs. Participants were required to indicate which character within each pairing they thought was the more responsible for each outcome. At the beginning of each trial, participants were presented with an outcome for two seconds. They were then presented with two characters sequentially (henceforth first and second characters). Participants were allowed to make a choice (choose the responsible character) from the onset of the second character.

**Figure 1:**
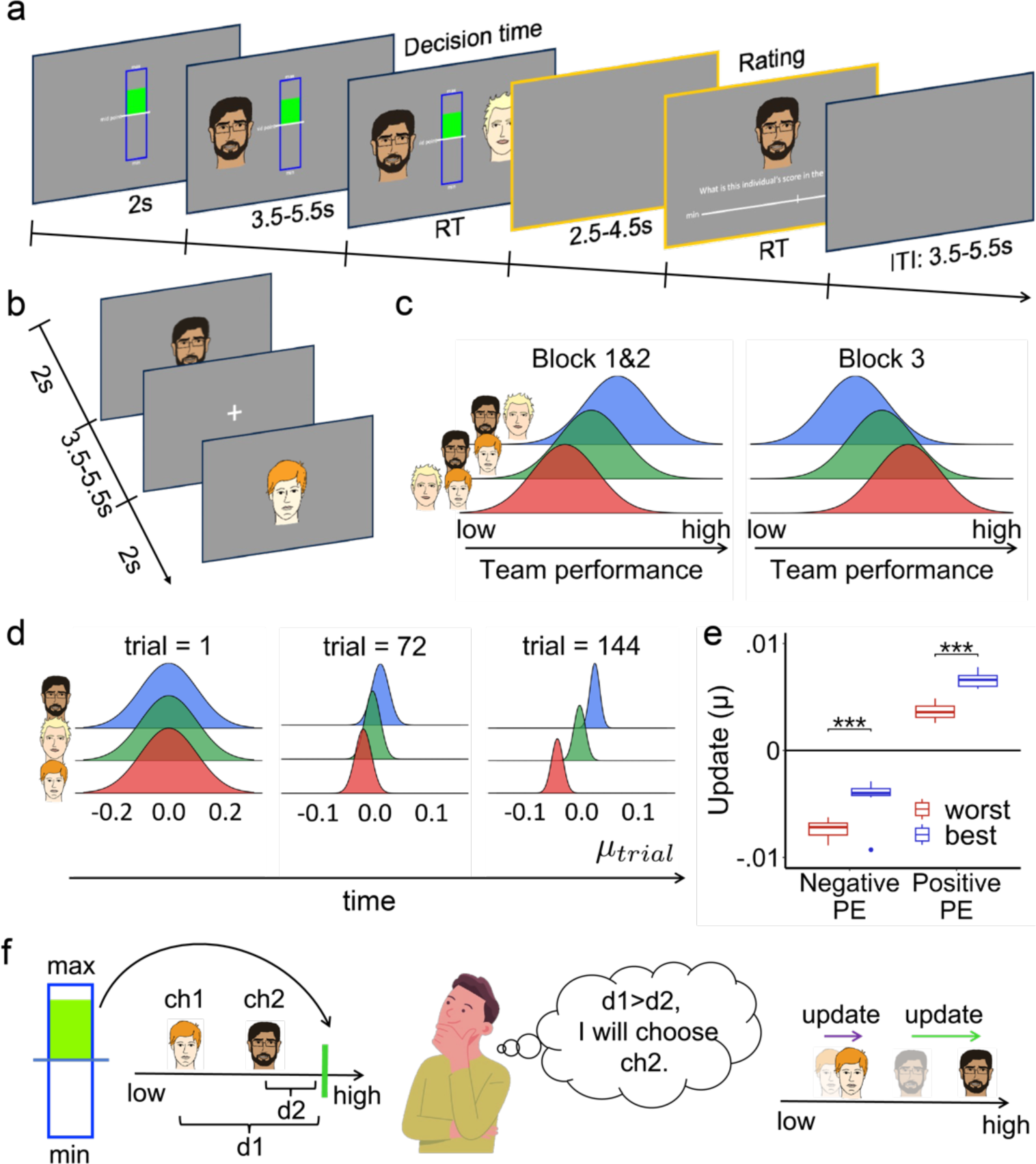
Experimental task and Simulation Results. **a** On each trial of the latent social cause inference task, participants, while undergoing fMRI, were sequentially presented with an outcome and two characters who had worked together to produce the outcome. Participants were required to indicate which character they thought was more responsible for the outcome. Participants were intermittently asked to rate the performance of one of the two characters, drawn at random too. **b** At the end of each session, participants were presented with the image of the characters in the task for 90 trials (30 trials per character) and were required to pay attention to the images. **c** The performances of each pair of characters resulted in three distinct but overlapping distributions. **d** The Bayesian learner tracked and updated the performance estimates for each character. The model learned the performance of the characters and became more certain of its estimate as the task progressed (compare the width of distributions between first and last trial). **e** The Bayesian learner updated its estimate of the better character in a pair more following a positive prediction error (PE) and updated its estimate of the worse character following negative PE. **f** Schematic of how a Bayesian learner or a human participant can solve the task. They can compute the unsigned |PE| for each character (a proxy for the probability that each character is responsible for the outcome). The character with a smaller unsigned |PE| is more likely to be responsible for the outcome (the outcome would be less surprising if it had been caused by them). They can then update their estimate of the responsible characters given the outcome, and to a lesser degree, their estimate of the character who was not chosen as the responsible character. In the boxplots in **e**, the black horizontal band inside the boxes indicate the median and the whiskers extend to the most extreme data samples that lie within the intervals spanning 1.5 times the inter-quartile range. \*\*\**p*<.001.

On each trial, the outcome for each pair of characters (three pairs in total) was generated from three distinct but overlapping distributions (See Methods). Participants could learn the abilities of the characters by paying attention to the outcomes that characters achieved in pairs (Figure 1c). One of the characters appeared in trials in which the outcome was drawn from the distribution with the largest or the middle mean (Figure 1c, henceforth the best character). Thus, this character has a high association with the outcome (a high ability to cause the outcome). One of the characters appeared in trials in which the outcome was drawn from the distribution with the smallest mean or middle mean (Figure 1c, henceforth the worst character). Thus, this character has a low association with the outcome (a low ability to cause the outcome). The other character appeared in trials where the outcomes were generated from the distributions with the highest or smallest mean. Thus, this character has a middling level of association with the outcome (a middling ability to cause the outcome). Participants completed three blocks of 72 trials each. In all three blocks, the task schedule (see Methods) was the same. However, in the third block, we introduced an ability reversal by swapping the worst and the best characters’ abilities (Figure 1c). At the end of some trials, one of the two characters was pseudorandomly selected, and participants were asked to rate the ability of the character on a rating scale ranging from “min” to “max”. After finishing the latent social cause inference task, participants performed a localiser task where they passively observed the images of the three characters for 90 trials (30 trials per each character, Methods, Figure 1b).

### Statistical procedure

We used the student t-test to assess significance in all regression models and Wilcoxon signed-ranked test for all other analyses. We applied Holm-Bonferroni Correction^19^ to correct for multiple comparisons, where applicable.

### Bayesian learning indicates asymmetric update

We first designed a Bayesian model to investigate how a normative model would solve this task (See Bayesian Learner in Methods). We ran the model on one of the trial sequences that we used for participants. On each trial we computed the posterior probability that a given outcome should be assigned to each character and assigned the outcome probabilistically to one of the characters (i.e., the chance of being selected was proportional to their posterior probability). The mean and variance associated with each character were also updated every time the character was held responsible for an outcome. We subsequently refer to these distributions as the estimates of ability for each character (Figure 1d). To visualise how the model assigned each outcome to each character, we computed a “prediction error” (PE: defined as the signed difference between the outcome and the mean of each of the two character’s posterior distributions at the start of the trial), and “updated” (defined as the subsequent change in the mean of the posterior distribution) following each outcome (see Methods). We plotted the update against PE separately for the best and the worst characters (Figure 1e) and found an asymmetry in the update: the update was larger for the better character in each pair following positive PEs. On the other hand, the update was larger for the worse character following negative PEs.

The Bayesian learner performs the task by implementing the following steps (Figure 1f): a) it tracks the characters’ abilities (their associations with the outcome), b) it computes the probability that each character is responsible for a given outcome using its estimate of characters’ abilities, c) it then chooses the character who is more likely to be responsible for the outcome (the character with a smaller unsigned PE, also referred to as, surprise, PE magnitude, or |PE|, and described in more detail below, is identified as responsible for the outcome because the outcome would be less surprising if it had been caused by them), and d) it finally updates its estimate of the ability of the responsible character given the outcome. These computations led to an asymmetric update in character abilities; the estimate for the character identified as responsible is updated more than the estimate for the character not identified as responsible. The intuition is that if the character is thought not to be responsible for the outcome, then the outcome should have little bearing on the participant’s view of that character. Importantly, however, while intuitive this implies a fundamental departure from all standard learning models where more learning (more UEA) occurs when outcomes are more surprising; here learning (UEA) is greater for characters who are more likely to be associated with such an outcome (with whom the outcome is more likely to have occurred). We therefore tested for signatures of such computations in how participants performed the task.

### Behavioural results

If participants perform the task in a manner reminiscent of the Bayesian learner, then we expect a negative relationship between the difference in unsigned PEs (the magnitude of the PE or |PE|) for the two characters and the probability of choosing a character. Such a pattern would suggest that participants chose characters as responsible for outcomes when it was less surprising that the character was the cause of the outcome. In addition, according to the Bayesian simulation, an outcome should have less impact on a character’s rating if the character’s unsigned PE (|PE|) is high (as the character is unlikely to have been responsible for that outcome). We thus moved on to test these predictions in our behavioural data. We first tested whether |PE| drives participants’ choice and second, whether the effect of outcome on rating is smaller when |PE| is high. To test the first prediction, we regressed participants’ choice against |PE| difference between characters (see LM1 in Methods) and found that difference in |PE| between the two characters had a significant impact on participants’ choice (*B*=–1.62, *SE*=0.13, *p*<.001; Figure 2a); participants chose the character with the smaller |PE|. To test the second prediction, we regressed participants’ rating of a character on a given trial against outcome of that trial, the unsigned |PE|, and the interaction between them (see LM2 in Methods). If the second prediction is correct, we expect a negative interaction between |PE| and outcome. This regression model showed that participants’ ratings of each character depended on the outcome they observed (*B*=0.51, *SE*=0.03, *p*<.001), but more importantly, there was a negative interaction effect between outcome and |PE| on rating (*B*=–0.32, *SE*=0.03, *p*<.001; Figure 2b).

**Figure 2:**
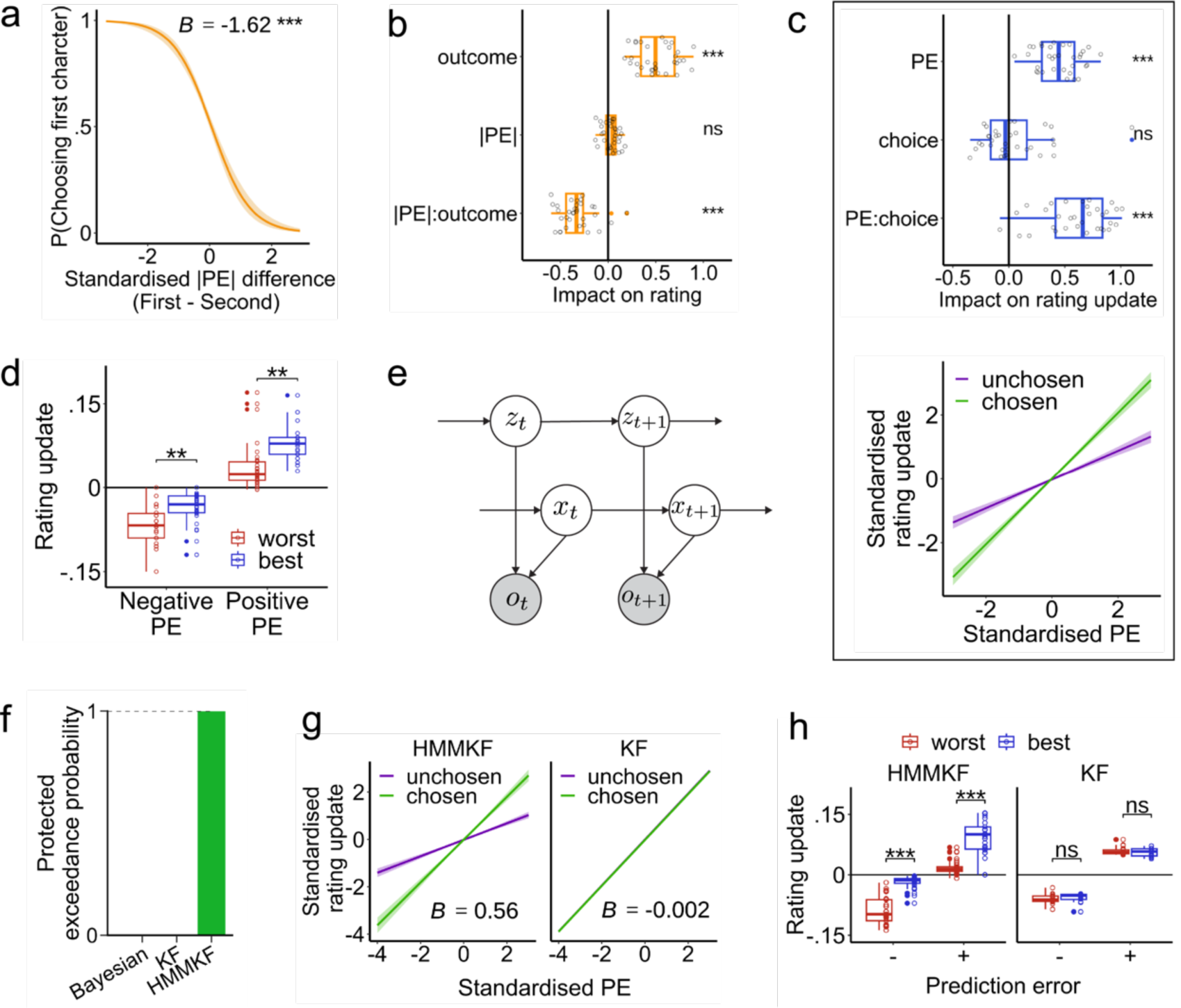
Behavioural and modelling results. **a** Probability of choosing the first character is plotted against the difference between unsigned prediction errors (PE magnitude or |PE|) for the two characters (first minus second character). Beta value was obtained using a regression model. **b** Beta values of a regression model performed for each subject separately in which rating was regressed onto outcome, |PE|, and their interaction. **c** top: Beta values of a regression model performed for each subject separately in which rating update (difference in a character’s rating in two consecutive time points that participant rated a character) was regressed onto PE, choice, and their interaction. Note that the participants’ choices indicate which character they held responsible for the outcome. bottom: rating update is plotted against PE separately for the chosen (responsible) and unchosen (non-responsible) characters. The update was larger for the chosen compared to the unchosen character. **d** Participants updated their estimate of the best character more following positive PEs, while their update of the worst character’s ability was larger following negative PEs. **e** Illustration of the generative process of the HMMKF model. The model assumes that the outcome on the current trial, *o_t_*, is noisily generated by one of the characters, encoded by the latent binary variable *z_t_*. The model further makes a Markovian assumption, whereby the current value of both variables depends on their value on the previous trial. **f** model frequency is plotted for all three models. Model frequency is zero for the Bayesian and KF models. **g-h** Unlike a generic KF model, the HMMKF model shows the key behavioural patterns that we observed. The update is larger for the responsible (chosen) character (g, left), pertaining to panel c top. It also shows the asymmetric update (**h**, left), pertaining to panel **d**. In **a**, **d**, **c** (bottom), **d**, **g**, and **f** error bars show standard error of the mean. In all boxplots, the black horizontal band inside the boxes indicate the median and the whiskers extend to the most extreme data samples that lie within the intervals spanning 1.5 times the inter-quartile range. Each dot indicates a beta value obtained from the regression model for each subject. The beta values were tested against zero using Wilcoxon signed-rank test. \*\*\**p*<.001, \*\**p*<.01, \**p*<.05, ns: not significant.

These results suggest that participants’ UEAs for each character were, indeed, less dependent on the observed outcome when |PE| was large, or in other words, when a character was unlikely to have been responsible for the outcome. To test this observation more directly, we regressed participants’ rating updates of each character against PE, responsibility (whether the character was held responsible for that outcome) as indexed by participants’ choices, and their interaction (see LM3 in Methods). Consistent with previous analyses, rating update was positively modulated by PE (*B*=0.45, *SE*=0.03, *p*<.001). However, unlike a standard reinforcement learning model, there was also a positive interaction between responsibility, as indexed by choice, and PE (*B*=0.58, *SE*=0.05, *p*<.001; Figure 2c, top), suggesting that a given PE had a larger impact on the rating update if the character was held responsible for that outcome (Figure 2c, bottom).

The results suggest that participants held the better (worse) character responsible for good (poor) outcomes. This in turn led them to update their rating of the better (worse) character more for good (poor) outcomes, leading to an asymmetry in UEA. To directly test this prediction, we computed rating updates for the best and the worst characters separately for positive and negative PEs (analogous to Figure 1e for the Bayesian learner). We found that for the first two blocks (before reversal), participants increased their ratings of the best character more than the worst character when the PE was positive (Figure 2d, *V*=67, *p*=.003, corrected for multiple comparisons), whereas they decreased their ratings of the worst character more than the best character following negative PEs (Figure 2d, *V*=61, *p*=.003; corrected for multiple comparisons). This observation is consistent with the simulation results using the Bayesian learner (Figure 1e).

Our results indicate that participants’ overall approach in performing the task is consistent with a Bayesian learner. However, it is not clear how well this model can capture participants’ trial-by-trial choice patterns. We thus designed two additional computational models and compared them to find the model which fitted participants’ behaviour better than the other models. The first model is a generic Kalman Filter (KF) which has been extensively used in neuroscience ^20–22^ to model learning and UEA. In addition, we designed a modified version of the KF by combining it with a hidden Markov model (henceforth HMMKF, Figure 2e). Unlike the KF model which updates both characters equally for a given outcome, HMMKF learns a latent state for each pair of characters and computes a responsibility value (R). This addition means that the KF component not only performs UEA but that the HMM component captures LCI. On each trial the HMMKF model only updates its estimate of the character identified as responsible for the outcome that is witnessed. We fit all three models to the participants’ trial-by-trial choices of the responsible character and compared the goodness of fit using protected exceedance probability^23,24^ (see Methods). The HMMKF model provided the best model fit (protected exceedance probability=1, Figure 2f, See Figure S1 for the summary of fitted parameters). For each participant, we inspected the choice and rating produced by the HMMKF and KF models using fitted parameters (obtained for each participant). Critically, the HMMKF model successfully reproduced the key behavioural patterns that we observed in participants’ data: the impact of responsibility on UEA and the asymmetric UEA (Figure 2g & 2h), while neither of these patterns was observed in the generic KF model. We therefore used the trial-by-trial estimate of choice, PE, and |PE| obtained from HMMKF model to analyse the fMRI data that we recorded during the task.

### The impact of context on LCI and UEA related behaviour

Finally, we sought to understand to what extent the results that we have reported so far depends on the specific context of our task. To investigate any potential effects of context on these computations, we first asked whether these computations are different in a non-social context. For this purpose, we employed a within subject design (experiment 2) where online participants completed a social and a non-social version of the task in randomised order. The social version was identical to experiment 1. The degree to which one might ask about responsibility in a completely non-social context might be contested but it is possible to ask about responsibility in the context of entities such as companies that are not social agents in the same way as individual people. In the non-social version, human characters were replaced by companies. Each company was assigned a fractal image. Participants were told that three companies collaborated to produce a product together and each outcome indicated the quality of their product. Every other aspect of the experiment (including trial sequence) was identical to the social version. We chose sample size (N=50) as we had fewer trials in this experiment compared to experiment 1.

We tested whether the negative interaction effect of outcome and |PE| on rating differed between the social and non-social contexts (LM2). We found that in both contexts there was a negative interaction between outcome and |PE| (social: *B=-.3, SE=.02, p<.001, nonsocial: B=-.3, SE=.02, p<.001*). Critically, there was not a three-way interaction between outcome, |PE|, and context (*B=-.01, SE=.02, p=.63*), suggesting that the effect was not different between the two contexts.

So far, in both experiments, participants were required to choose the responsible character on each trial. It might be argued that the interaction effect between |PE| and outcome on rating, a critical marker of an inference process, is prompted by the forced contemplation and explicit singling out of one character as responsible. To test whether this task feature might have led the participants to only update the responsible character, we conducted experiment 3 in which participants (N=50) performed a task in which they were not required to identify the responsible character in either social or non-social situations. All other aspects of this experiment were identical to experiment 2. We found that in both social and nonsocial contexts there was still a significant negative interaction effect between outcome and |PE| (social: *B=-.09, SE=.02, p<.001, nonsocial: B=-.08, SE=.02, p<.001*). Again, there was no difference between the two contexts (*B=.002, SE=.02, p=.93*).

The results of experiments 2 and 3 indicate that the LCI and UEA related computation that we report here reflect a general computational strategy that humans use regardless of specific task context or task features. Inferring responsibility occurs naturally, even when the participants are not explicitly required to identify the responsible character.

### Neuroimaging results

#### Medial prefrontal cortex involvement in updating characters’ abilities

Having established participant behaviour, we turned to fMRI to investigate how the human brain implemented the computations pertinent to this approach. We first turned our attention to UEA but in the next section below we sought the neural mechanism mediating the LCI process on which UEA depended. We performed a whole-brain computational model-based general linear model (GLM) using the PE of the first character as the parametric modulator of BOLD activity at the time of first character presentation (Methods, GLM1). We found three clusters which were negatively modulated by the PE of the first character, dorsomedial prefrontal cortex (dmPFC), ventromedial prefrontal cortex (vmPFC), and left orbitofrontal cortex/anterior insular cortex (OFC/AI; Figure 3a; see Table S1 for full results of GLM1).

**Figure 3:**
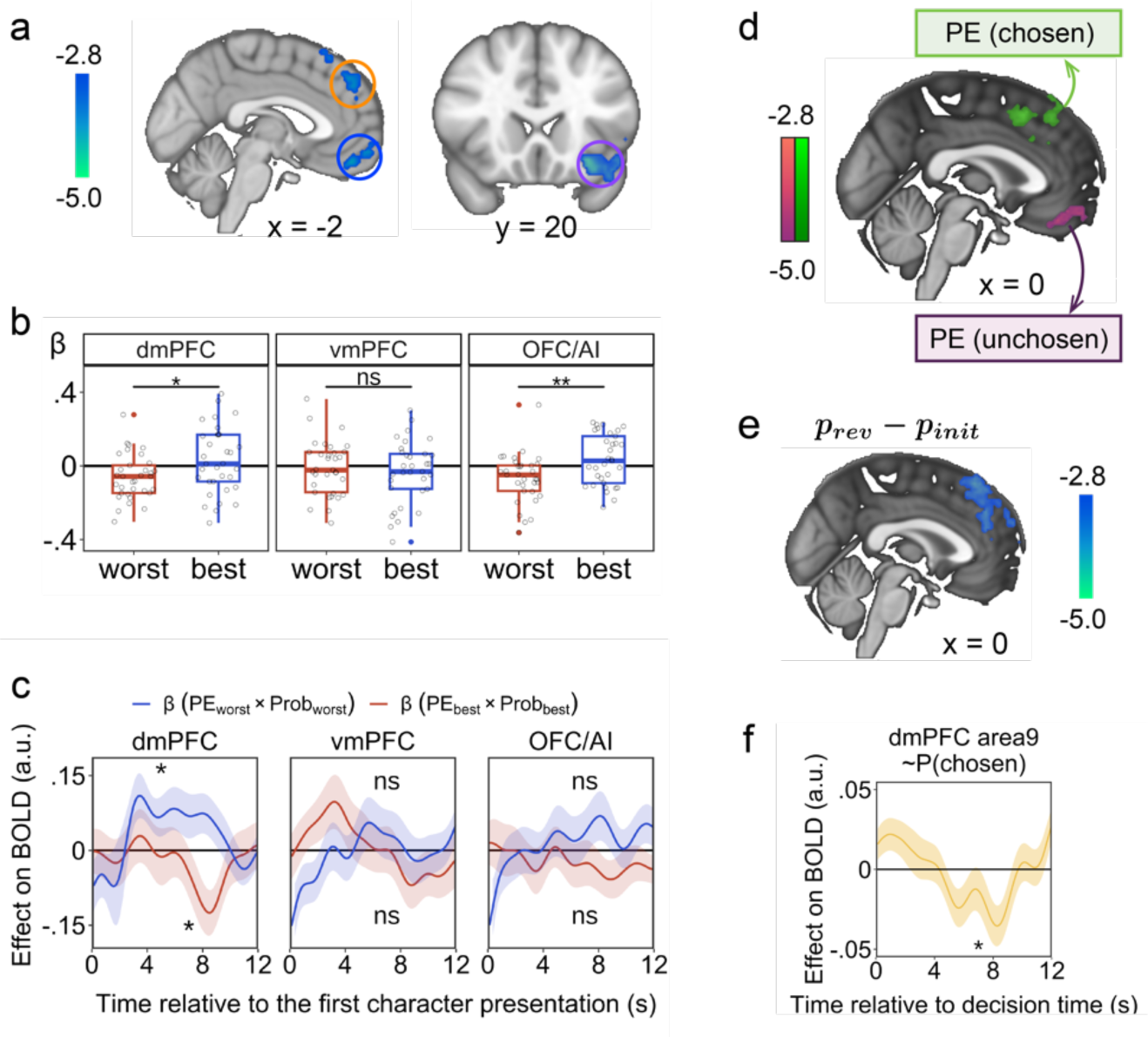
dmPFC contribution in the task. **a** Whole-brain analysis revealed three brain areas which encoded prediction error (PE) at first character presentation, including dmPFC (peak Z=-3.65, MNI: x=-4, y=48, z=36), vmPFC (peak Z=-3.86, MNI: x=-4, y=60, z=-8), and OFC/AI (peak Z=-4.24, MNI: x=-30, y=20, z=-12). **b** At first character presentation, encoding of PE was different for the best and the worst characters in dmPFC and OFC/AI. **c** A regression model indicates that, at first character presentation, BOLD activity in dmPFC but not vmPFC or OFC/AI, is correlated with the interaction between PE and the probability that the first character is responsible for the outcome (*p_init_*); this interaction is a critical determinant of rating update (Figure 2c, top). The interaction term is encoded with different signs for the best and the worst characters (positive and negative for the best and worst characters, respectively). **d** At the time of choice, dmPFC and vmPFC encoded PE for the chosen and unchosen characters, respectively (dmPFC cluster: peak Z=-3.95, MNI: x=8, y=18, z=50; vmPFC cluster: peak Z=-3.80, MNI: x=0, y=40, z=-20). **e** The change in the estimate of responsibility for the first character – the difference between the probability that the first character is responsible after seeing just the first character and then after also seeing the second character on a trial (*p_rev_* − *p_init_*) is represented in dmPFC (peak Z=-3.93, MNI: x=-2, y=34, z=54). **f** dmPFC also encodes the probability that the character finally chosen is responsible for the outcome. In the boxplots in **b**, the black horizontal band inside the boxes indicate the median and the whiskers extend to the most extreme data samples that lie within the intervals spanning 1.5 times the inter-quartile range. In c and f, the lines and shaded areas show the mean and standard error of the mean (s.e.m), respectively. \*\*\**p*<.001, \*\**p*<.01, \**p*<.05, ns: not significant.

However, both behavioural and computational modelling demonstrated asymmetry in UEA (Figure 2d) suggesting that the brain holds a similarly asymmetric representation of PE. Such an asymmetric representation can be quantified by comparing the representation of PE between the best and the worst characters. With this aim, we extracted time course activities within these regions (Methods, TC1.1) and compared the response of these areas to PE separately for the best and the worst characters. Intriguingly, this analysis revealed that the responses of dmPFC and AI/OFC were significantly different between the worst and the best characters (more positive response to PE of the best character; Figure 3b, dmPFC: *V*=420, *p*=.023; OFC/AI: *V*=443, *p*=.009, vmPFC; *V*=256, *p*=.672, corrected for multiple comparisons). This asymmetric representation of PE could have arisen because these areas did not simply encode PE but did so only when the probability that the first character was responsible for the outcome was high. In another words, these areas encoded the asymmetric UEA. Just as in behaviour, where the asymmetry in UEA can be traced to the interaction between PE and the initial probability that the character is responsible for the outcome, the same is true in the neural data (*p_init_*, as opposed to the revised probability estimate, *p_rev_*, reached after observing the second character; see Methods).

To test this hypothesis, we extracted the regions’ activity time courses and regressed them against probability, PE (both obtained from our HMMKF model), and their interaction, separately for the best and the worst characters (see Methods, TC1.2). Intriguingly, we found a positive interaction effect of probability and PE on dmPFC BOLD for the best character (Figure 3c, *W*=376, *p*=.045, corrected for multiple comparisons), and a negative effect of probability and PE on dmPFC BOLD for the worst character (Figure 3c, *W*=159, *p*=.031, corrected for multiple comparisons). These results mean that the activity of dmPFC was highest when there was a positive PE for the best character (when UEA is high) and there was a negative PE for the worst character (when again UEA is high); the asymmetry in dmPFC UEA (Figure 3c,d) resembles the asymmetry in UEA in behaviour (Figure 2d). We did not observe the same pattern in vmPFC or OFC/AI (Figure 3c, vmPFC (best/worst): *W*=195, *p*=.129/*W*=291, *p*=.429; OFC/AI (best/worst): *W*=286, *p*=.543/*W*=293, *p*=.761, corrected for multiple comparisons). The close analogy between dmPFC activity and behavioural update suggests that dmPFC updates the characters’ abilities after inferring the latent causes (LCI). We test this idea further in experiment 4 below.

The asymmetric UEA (Figure 2c) might suggest the brain kept a separate representation of the PE of the chosen and unchosen characters at decision time. This raises the possibility that at the time of the choice, when the update happens, the brain represents the PE of both chosen and unchosen characters. We thus conducted a whole-brain GLM (GLM2, Methods) and found that PE of the chosen character was represented in dmPFC while PE of the unchosen character was represented in a more ventral portion of mPFC, in an area referred to as ventromedial prefrontal cortex (vmPFC) (Figure 3d; see Table S2 for all significant clusters).

The initial probability that the first character is responsible for the outcome (*p_init_*; see Methods) was calculated under the assumption that the two other remaining characters were equally likely to appear as the second character (See Eq(21) in Methods). However, when the second character appeared, this probability estimate for the first character could have been revised (*p_rev_*) in light of the new information furnished by the identity of the second character (See Eq(22) in Methods). We thus asked whether the brain revised its initial estimate that the first character was responsible for the outcome and calculated the revised probability (*p_rev_*). We therefore ran a new whole-brain GLM (GLM3) and tested for the neural correlates of the difference between the revised and the initial probability (*p_rev_* − *p_init_*). Intriguingly, we found that this information was again encoded in dmPFC (Figure 3e; see Table S3 for full results of GLM3). This result indicates that at the time of the second character, the dmPFC revised its previously held estimate that the first character was responsible for the outcome (UEA), suggesting that dmPFC might represent the probability that the final chosen character was responsible for the outcome (*p_chosen_*). To test this prediction, we extracted the time course of an anatomical mask of dmPFC encompassing area 9^25^ and found that this area, at the time of second character presentation (or decision time), indeed encoded *p_rev_* for the character finally chosen as responsible (*W*=143, *p*=.015, corrected for multiple comparisons; Figure 3f). We note that we used a mask derived from a previous anatomical study to avoid any selection bias in determining the dmPFC region analysed (Figure 3a and 3d). However, we found the same results when we repeated this analysis on our previous dmPFC ROI which encoded the PE of the first character (Figure S2). Finally, using a whole-brain GLM (Figure S3) we report brain areas which encoded a decision signal (i.e., the probability that the chosen character was responsible for the outcome minus this probability for the unchosen character) at the time of choice. In addition, we show elevated functional coupling between areas that encoded the decision signal and dmPFC (as a function of PE of the *chosen* character) and vmPFC (as a function of PE of the *unchosen* character; Figure S4).

So far, our results indicate that dmPFC plays a crucial role in UEA: it encodes PE for the first character at first character presentation. The PE signal in dmPFC is only present when the first character is likely to be responsible (i.e., *p_init_* is high). As described in Figure 1f, asymmetric UEA is dependent on first performing LCI. We therefore turned to more sophisticated multivariate pattern analysis of our fMRI data to understand how the brain achieves LCI.

#### Hippocampus involvement in inferring latent cause

The hippocampus has been linked to constructing a representation of physical space and the space of states that describe a possible task^13^. In addition, the hippocampus is suggested to construct a predictive map of future state occupancies^26^ given the current state estimate. Arguably in our task estimating the current state and predicting the next is equivalent to finding the character most likely to be responsible for the outcome witnessed on each trial; identifying the character responsible for the outcome is what must be achieved by LCI when each outcome is observed. Given that experiment 3 had demonstrated the automaticity of the inference process regardless of the need for an explicit responsibility judgement, we hypothesized that the brain initiates the inference process at the time of outcome, even before seeing any characters. To test this hypothesis, we applied a decoding approach (See Decoding Analysis in Methods). With this aim, we trained a support vector machine (SVM) classifier on the localizer data that we obtained from each participant^27^ (Figure 1b). Intriguingly, we found that a cluster in left hippocampus was the only brain region in which we could significantly decode the most likely character (i.e., identify the latent cause of the outcome), prior to any character (the potential cause) presentation, above chance level after correcting for multiple comparisons (Figure 4a). By contrast, when participants were not identifying characters by LCI but were instead directly observing them on a screen, then character identity was no longer decodable from hippocampal activity, but instead only from fusiform face area (FFA; Figure 4a and Figure S5). Therefore, unlike FFA, hippocampal activity does not reflect perception of character identity but rather inference of identity. There was no confound between brain activity when character identities were inferred from outcomes, and the next character identities directly observable on the screen; the character inferred at outcome and the next character to appear on the screen were the same in only a small proportion of trials (Figure S6).

**Figure 4:**
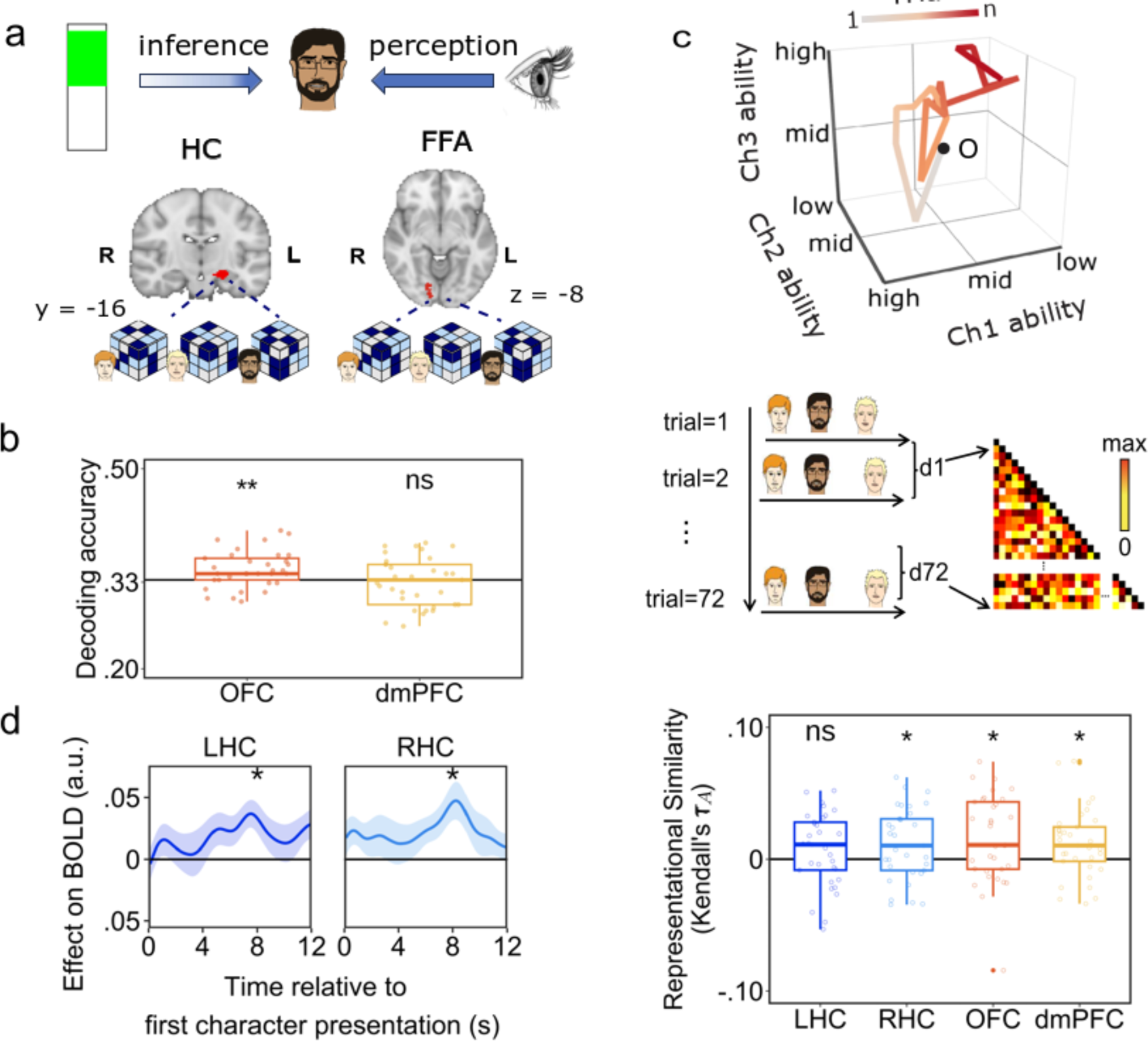
Hippocampus contribution in our task. **a** left, Whole-brain searchlight classification revealed that the most likely character to have caused a given outcome could have been decoded from activity in a cluster in left hippocampus (LHC, *Z*=2.48; voxel-wise threshold at *p*<.001, cluster correction at *p*<.05) at the time of LCI when the outcome was presented and before any characters were seen on that trial. Right, However, at the time of the presentation of the first character, the identity of the character was decodable from activity pattern of FFA, but not hippocampus. **b** ROI-based MVPA analysis showed the most likely character could be decoded from OFC at outcome time, but not in dmPFC. **c** Top: we asked whether activity pattern in the hippocampus represents the 3-dimensional ability space that participants traverse during the task. The plot shows evolution of the ability space for an example participant. Middle: We constructed our model RDM based on the trial-by-trial change in all three characters’ abilities in the task. Bottom: Representational similarity (correlation between model and brain RDMs) is plotted for four brain regions. **d** The probability that the first character is responsible for the outcome (*p_init_*) is encoded in RHC and LHC (top panel), but not in OFC or dmPFC (bottom panel), activity. In the boxplots in **b** and **c**, the black horizontal band inside the boxes indicate the median and the whiskers extend to the most extreme data samples that lie within the intervals spanning 1.5 times the inter-quartile range. Each dot indicates a participant. In **d**, error bars indicate standard error of the mean (s.e.m). \*\*\**p*<.001, \*\**p*<.01, \**p*<.05, ns: not significant.

Finally, we determined decoding accuracy of character perception in the hippocampus ROI. Using Bayesian statistics^28^, we found significant evidence in favour of the null as quantified using Bayes factor (*BF*_10_=.2, W=249). We then used Exemplar Dissimilarity Index (EDI) to investigate whether there was any information about character identity in hippocampus during perception (first character presentation). The EDI asks whether a multivariate pattern is more stable across scan runs within conditions compared to between conditions^29^. Condition was defined as the identity of each character (See Methods). We found that the hippocampus does not carry any information about character identity during perception at the time of first character presentation (LHC, p=.96, W=177, *BF*_10_=.07*, RHC:* p=.48, W=315, *BF*_10_=.55, corrected for multiple comparison). However, on repeating this analysis in dmPFC, we found that the activity pattern in this area carries information about the identity of the characters (W=375, p=.04) at the time of first character presentation.

We then performed two ROI-based decoding analyses. First, if dmPFC is only involved in the UEA process, we expect that it might not code for the responsible character at the time of outcome. Such a finding would suggest that dmPFC is not involved in LCI related computation. Decoding accuracy in dmPFC was below chance level, even when no whole-brain correction was applied (dmPFC: *W*=252, *p*=.822; Figure 4b, corrected for multiple comparison). We used Bayesian statistics^28^ in a dmPFC ROI analysis and found significant evidence in favor of the null as quantified using Bayes factors (*BF*_10_=.21, W=290, Figure 4b). Second, since the orbitofrontal cortex (OFC) has been implicated in representing learned and hidden task states^4,30,31^, we conducted another ROI-based analysis focusing on the decoding accuracy of OFC. This analysis revealed significant evidence that OFC encodes the most likely character (W=396, p=.011, corrected for multiple comparison, Figure 4b). These results suggest clear dissociation between hippocampus and dmPFC in performing LCI and UEA related computations, respectively.

Representing the most likely character is only possible through knowledge of the characters’ abilities (the participant’s estimates of association between a character and outcome). We thus asked whether the activity pattern within hippocampus carried a map-like 3-dimensional (3D) representation of characters’ abilities (Figure 4c, top). With this aim, we used representational similarity analysis (RSA)^32,33^ and asked whether the distances between all three characters’ abilities across trials were correlated with the hippocampus activity pattern. Our model representational dissimilarity matrix (RDM) was thus constructed using trial-by-trial distances between the abilities of all three characters. With this aim, we first computed the model-derived ability estimates of all characters for each trial (a vector with three entries for each trial). The model RDMs were constructed by computing the Euclidian distance between these vectors (Figure 4c, middle). Consistent with our prediction, the activity pattern of hippocampus was correlated with the estimate of the characters’ abilities (Figure 4c, bottom, LHC: *W*=366, *p*=.065; RHC: *W*=398, *p*=.035, corrected for multiple comparisons). Therefore, instead of representing the participant’s position in 2- or 3-D physical space as occurs in a spatial navigation task^34^, or a cognitive map of abstract value space^35^, in this task, hippocampus represents an abstract map of character ability space (Figure 4c). On repeating this analysis on activity patterns extracted from OFC and dmPFC we found both of these areas also represented character ability spaces (Figure 4c, bottom, dmPFC: *W*=413, *p*=.025; OFC: *W*=421, *p*=.022, corrected for multiple comparisons). The representation of character ability in dmPFC is consistent with our univariate results which demonstrated representation and potential update of characters’ PE at first character presentation and decision time (Figure 3). Therefore, while the dmPFC and hippocampus both maintain representations of relative character abilities, the representation of the most likely character, when it is obtained by LCI on presentation of a new outcome, can only be decoded from hippocampus but not dmPFC.

We next reasoned that if hippocampus achieves LCI of the most likely character before presentation of any of the characters, it might also compute the probability that the first character was responsible for the outcome (*p_init_*) at first character presentation. To test this prediction, we regressed hippocampus time course activity against *p_init_*, PE, and their interaction. Consistent with our prediction, hippocampus activity was modulated by *p_init_* (Figure 4d, LHC: *W*=425, *p*=.030; RHC: *W*=427, *p*=.027; corrected for multiple comparisons), while neither of the two other regressors had significant effects on its activity (Figure S7). OFC and dmPFC activity were not modulated by *p_init_* (Figure 4d, OFC: *W*=219, *p*=1.000; dmPFC: *W*=293, *p*=1.000; corrected for multiple comparisons). So, although dmPFC activity clearly reflects changes and revisions in ability estimates (for example, it represents the difference between the revised and the initial probability: *p_rev_* − *p_init_*), its activity does not so clearly reflect maintenance of that estimate, *p_init_*. In summary, while the character ability space was represented in both areas, the UEA signal was prominent in dmPFC, suggesting a close relationship between its activity and updating, while LCI-related activity was prominent in hippocampus.

#### dmPFC-hippocampus interaction for solving the performing the inference and the update processes

So far, our results suggest that hippocampus and dmPFC perform LCI and LCI-dependent UEA respectively, but these different specializations are related to some activity patterns common to both areas. We therefore hypothesized that the interaction between these two areas was necessary for performing the entire process of LCI and UEA. We thus predicted that at first character presentation, functional coupling between hippocampus and dmPFC should increase as a function of the strength of association between the character and outcome. To test this prediction, we conducted a psychophysiological interaction (PPI) analysis by regressing dmPFC BOLD against the interaction between hippocampal BOLD (as the physiological variable) and *p_init_* (as the psychological variable), while controlling for main effects of both variables (See PPI analysis in Methods). We found that, indeed, functional connectivity between dmPFC and left hippocampus increased as a function of *p_init_* (Figure 5a, left hippocampus: *W*=154, *p*=.024; right hippocampus: *W*=269, *p*=.844, corrected for multiple comparisons).

**Figure 5:**
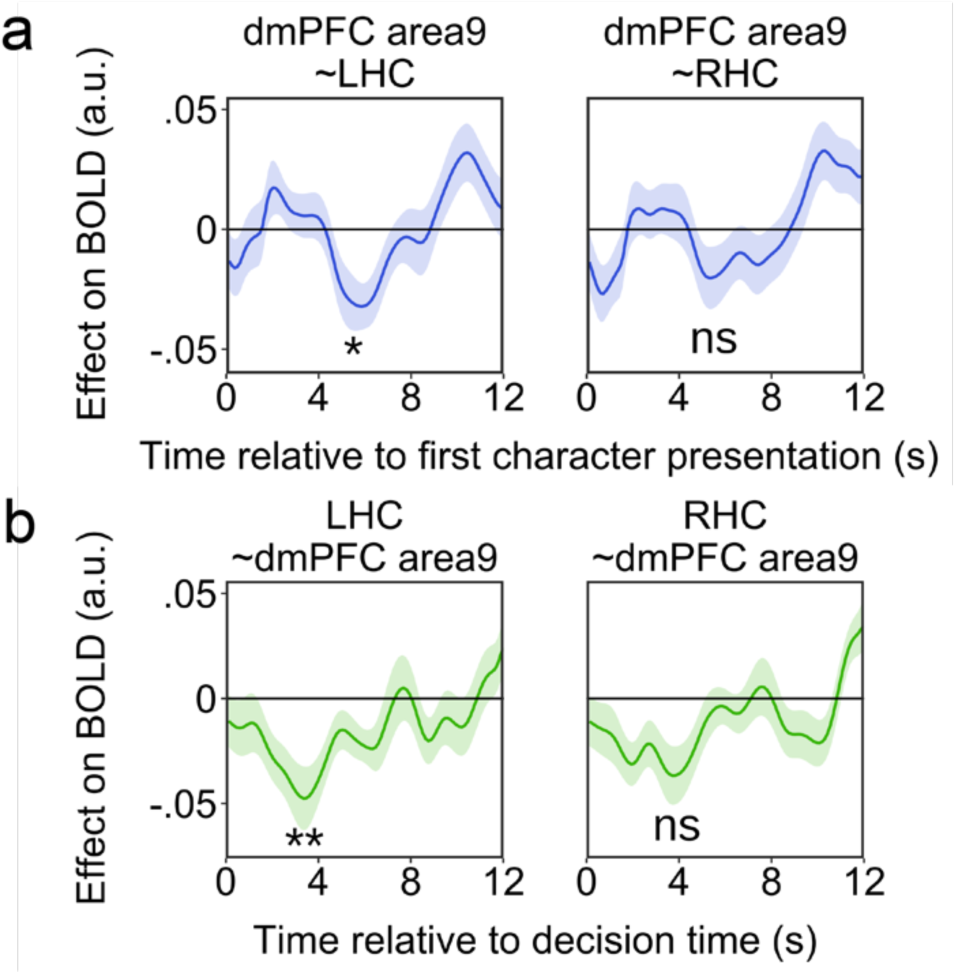
dmPFC-hippocampus interaction. **a** At the time of first character presentation, functional connectivity between left hippocampus and dmPFC (area9) increases as a function of the probability that the first character is responsible for the outcome. **b** At the time of choice (decision time), functional connectivity between left hippocampus and dmPFC area9 increases as a function of the rating update of the chosen character. The lines and shaded areas show the mean and standard error of the mean (s.e.m), respectively. \*\**p*<.01, \**p*<.05, ns not significant.

We also asked, whether functional coupling between these two areas increased, as a function of the size of UEA for the chosen character at decision time. With this aim, we performed a second PPI analysis by regressing hippocampal BOLD against the interaction between dmPFC BOLD (as a physiological variable) and the magnitude of the update signal for the chosen character (as the psychological variable) while controlling for the main effects. Again, we found that the functional coupling between these areas increased as a function of the size of the update signal (Figure 5b, left, *W*=135, *p*=.010; right *W*=171, *p*=.052, corrected for multiple comparisons).

#### Experiment 4: causal manipulation of dmPFC impairs asymmetric update

To establish the specificity and causal necessity of the dmPFC to UEA rather than LCI, we ran a within-subject experiment (n=19) where we applied continuous theta-burst stimulation (cTBS) to either dmPFC or a control area (vertex) or sham (where no stimulation was applied) during separate visits to the laboratory (Figure 6a). In each session of the experiment, participants completed three blocks of the task (using a version almost identical to the fMRI experiment 1; see Methods). In dmPFC or vertex sessions, cTBS was applied immediately before starting the task. The sham session was included so that we could disentangle any cognitive impairment following dmPFC cTBS from any potential impairment following active cTBS to any brain area (e.g., vertex). Here we focus on the comparison between dmPFC and vertex cTBS. In Figure S8, by comparing vertex versus sham, we show that cTBS over vertex did not lead to the kind of impairment that we observed following dmPFC cTBS that we report here. The effect of cTBS peaks 10 minutes after stimulation and disappears after 30 minutes^36^. We therefore focused our analysis on the first two blocks which were completed in the first 25 minutes after applying cTBS.

**Figure 6:**
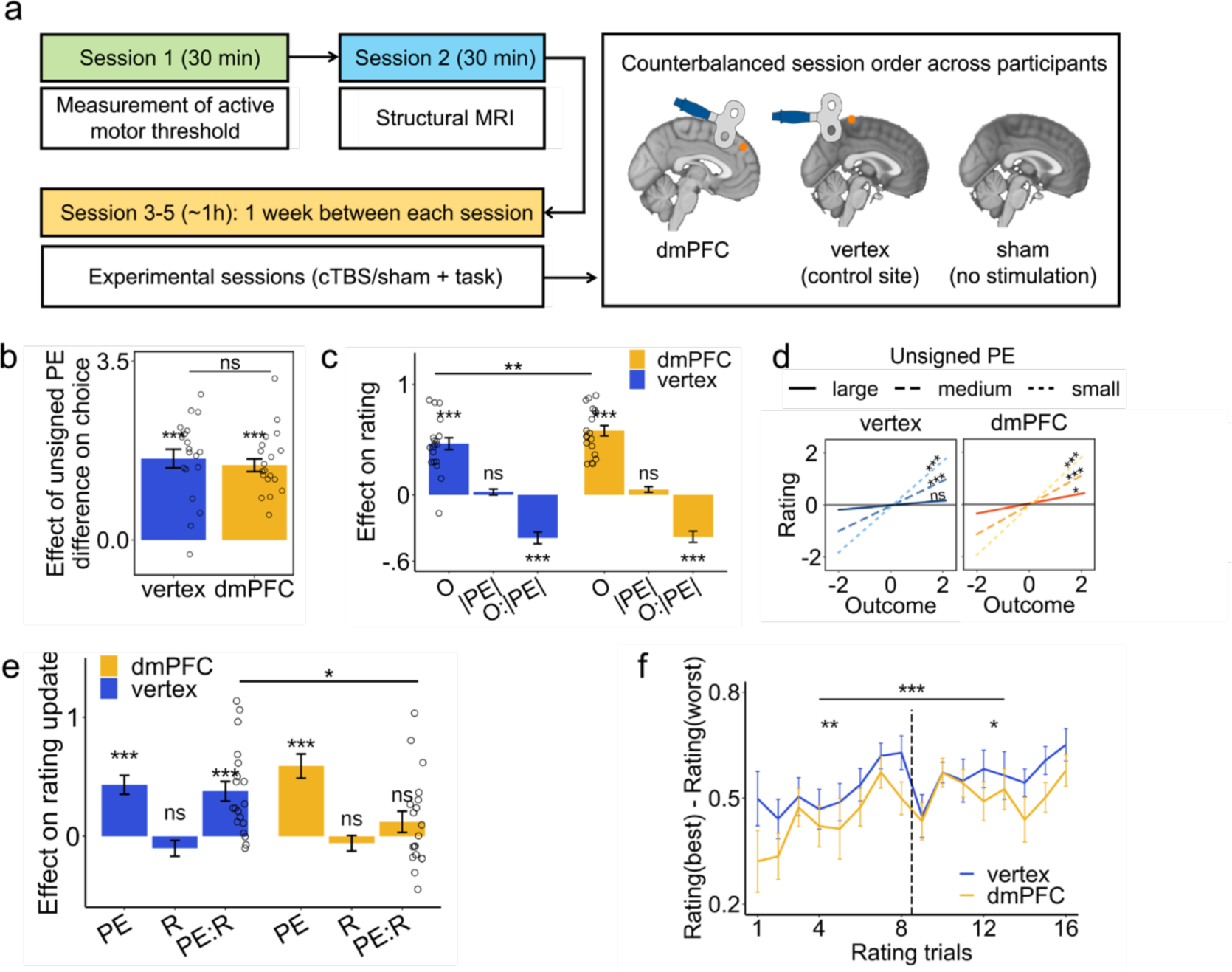
Disruption of dmPFC. **a** cTBS procedure. Each participant attended five sessions in total. For each participant, we measured their active motor threshold in Session 1, obtained a structural MRI scan in Session 2, and applied cTBS either to dmPFC or a control area (vertex) or sham (where no stimulation was applied) in subsequent three experimental sessions (Session 3-5) in pseudo-randomised order. **b** The effect of unsigned PE on choice was quantified using a multilevel regression model. This effect was not different between dmPFC and vertex stimulation conditions. **c** Using a multilevel regression model, we quantified the effect of unsigned PE, outcome, and their interaction on rating. We found that outcome had a higher impact on rating, in dmPFC compared to vertex condition. **d** The effect of outcome on rating is plotted for large, medium and small unsigned PEs. While the effect of outcome on rating is not significant for large PEs in vertex condition, in dmPFC condition this effect is significant. **e** Using a multilevel regression model, we quantified the effect of PE, responsibility (R), and their interaction on rating update. We found that dmPFC cTBS significantly decreased the effect of the interaction between PE and responsibility. **f** Rating difference between the best and the worst characters was lower in dmPFC compared to vertex condition. In **b**, **c**, and **d**, each point represents the beta value obtained from the regression model for each subject. In **f**, the error bars represent standard error of the mean (SEM). See detailed cTBS results (including regression coefficients and comparisons between conditions) in Table S6. \*\*\**p*<.001, \*\**p*<.01, \**p*<.05, ns not significant.

We first analysed participants’ choice behaviour and compared it across the two conditions. Our fMRI data indicated that participants’ choice of responsible character (latent cause) was guided by the difference between the characters’ unsigned PE (|PE|): a character with a larger |PE| was less likely to be chosen. If dmPFC cTBS impairs this aspect of inference, it suggests a causal role for dmPFC in LCI. To investigate this possibility, we regressed participants’ choice against the difference between the characters’ unsigned |PE| (the same as LM1; Figure 6b). We found that participants’ choices were guided by |PE| in both conditions (vertex: *B*=1.59, *SE*=0.18, *p*<.001; dmPFC: *B*=1.47, *SE*=0.12, *p*<.001), with no difference between dmPFC and vertex stimulation (*V*=116, *p*=.418). Bayesian statistics revealed moderate evidence in favour of the null hypothesis, suggesting that this lack of effect is evidence of absence rather than an absence of evidence (*BF*_10_ = .3). Therefore, dmPFC stimulation did not have any impact on participants’ ability to achieve LCI.

We then tested whether dmPFC stimulation had any impact on UEA. For this purpose, we first investigated the effect of |PE|, outcome, and their interaction on rating of each character (the same as LM2). For both vertex and dmPFC stimulation conditions, we observed a positive effect of outcome (vertex: *B*=0.48, *SE*=0.05, *p*<.001; dmPFC: *B*=0.56, *SE*=0.05, *p*<.001) and a negative interaction effect (vertex: *B*=-0.39, *SE*=0.05, *p*<.001; dmPFC: *B*=-0.37, *SE*=0.05, *p*<.001) on rating. However, participants’ ratings were influenced more by the current outcome in the dmPFC compared to vertex condition (*V*=23, *p*=.002; Figure 6c), while the interaction effect was not different between the two conditions (*V*=91, *p*=.891). The negative interaction effect indicates that an outcome has less impact on the UEA for a character’s when the character’s unsigned |PE| is very high (Figure 6c); in other words, the outcome has less impact on changing one’s estimation of a character’s ability when the outcome is surprising. The elevated effect of outcome on rating following dmPFC cTBS suggests that this is less so the case after dmPFC disruption; outcomes had a high impact on UEA even when the unsigned |PE| was high in the dmPFC condition. To test this interpretation, we performed a Simple Slopes Analysis^37^ and tested the effect of outcome on rating in trials with low, medium, and high |PE| separately. In both vertex and dmPFC conditions, we found that in the small and medium |PE| trials, outcome had a significant effect on rating (see statistics for small and medium |PE| in Table S5). However, and confirming our interpretation, in large |PE| trials, while outcome did not have any impact on rating in the vertex condition (*B*=0.09, *SE*=0.07, *p*=.241; Figure 6d), it had a significant impact on rating in the dmPFC condition (*B*=0.19, *SE*=0.08, *p*=.032; Figure 6d; see statistics for small and medium |PE| in Table S5).

These findings suggest that disruption of dmPFC activity impaired UEA dependent on LCI but not the LCI process itself. To confirm this interpretation using a different analysis, we tested the effect of PE, responsibility, and their interaction on rating update (same as LM3; Figure 6e). The rating update was significantly driven by PE in both stimulation conditions (vertex: *B*=0.44, *SE*=0.08, *p*<.001; dmPFC: *B*=0.58, *SE*=0.10, *p*<.001). However, there was only a positive interaction effect between PE and responsibility on rating update in the vertex stimulation condition (vertex *B*=0.38, *SE*=0.08, *p*<.001, dmPFC *B*=0.12, *SE*=0.09, *p*=.189) and the effect of the interaction term was significantly lower in the dmPFC versus vertex condition (*V*=148, *p*=.032; Figure 6e).

In summary, inferred responsibility (resulting from LCI) has less impact on UEA following dmPFC cTBS. Consequently, it should become more difficult for participants to distinguish between the best and the worst characters. To examine this possibility, we computed the difference in participants’ ratings of the best and the worst characters and compared this difference across the two conditions. Using a multilevel model, we found that the difference in ratings between the best and the worst characters was significantly smaller in the dmPFC compared to the vertex condition (Figure 6f; Block 1: *B*=–0.08, *SE*=0.03, *p*=.005, Block 2: *B*=– 0.05, *SE*=0.02, *p*=.014; aggregated: *B*=–0.07, *SE*=0.02, *p*<.001, corrected for multiple comparisons).

## Discussion

This study investigated an important social problem – how we work out which individual is responsible for which outcome when individuals work together in teams. However, the heart of this problem concerns a fundamental question common to many cognitive domains: when the cause of an outcome cannot be directly observed but must be inferred, how does the brain solve the interrelated problem of discovering the latent cause of this outcome (LCI) before updating the estimate of the strength of association between the outcome and cause (UEA) which, in the task’s social setting, corresponds to updating the estimate of the character’s ability. When we observe an outcome which is achieved by a group of people, the extent to which we update our estimate of their ability might not be equal for all members of the group. Instead, we update our estimates of their abilities according to how responsible we consider them to be for the outcome as a result of LCI.

We identified two brain regions which played distinct but complementary roles in LCI and UEA contingent on LCI (Figure 7). Our results suggest that the hippocampus plays a key role in LCI. We found that when participants were only presented with the outcome, the hippocampus already represented the most likely character to have caused it (Figure 4a). In other words, when the characters were directly perceived, their identities were encoded in visual association cortex (FFA; Figure S5) but when their identities were inferred by LCI from outcomes for which they might be responsible, their identities were encoded in hippocampus and OFC. Performing this inference process was only possible because the hippocampus kept a 3-dimesnional map of characters’ ability space, quantified by RSA (Figure 4c). Rather than representing a physical space, hippocampus represents participants’ positions in an abstract task space in terms of estimates of the abilities of each of the three characters. Consistent with these observations, when the first character appeared on the screen, the hippocampus encoded the probability that the first character was responsible for the outcome (Figure 4d). These observations suggest that the hippocampus performs LCI. This suggestion is broadly consistent with a proposed role for the hippocampus in discovering and constructing state space, especially when it is hidden^2,34,38,39^. The social nature of the task, in which multiple characters interact in different ways on different occasions, provides an ideal context for LCI investigation that is intuitive for participants.

**Figure 7.**
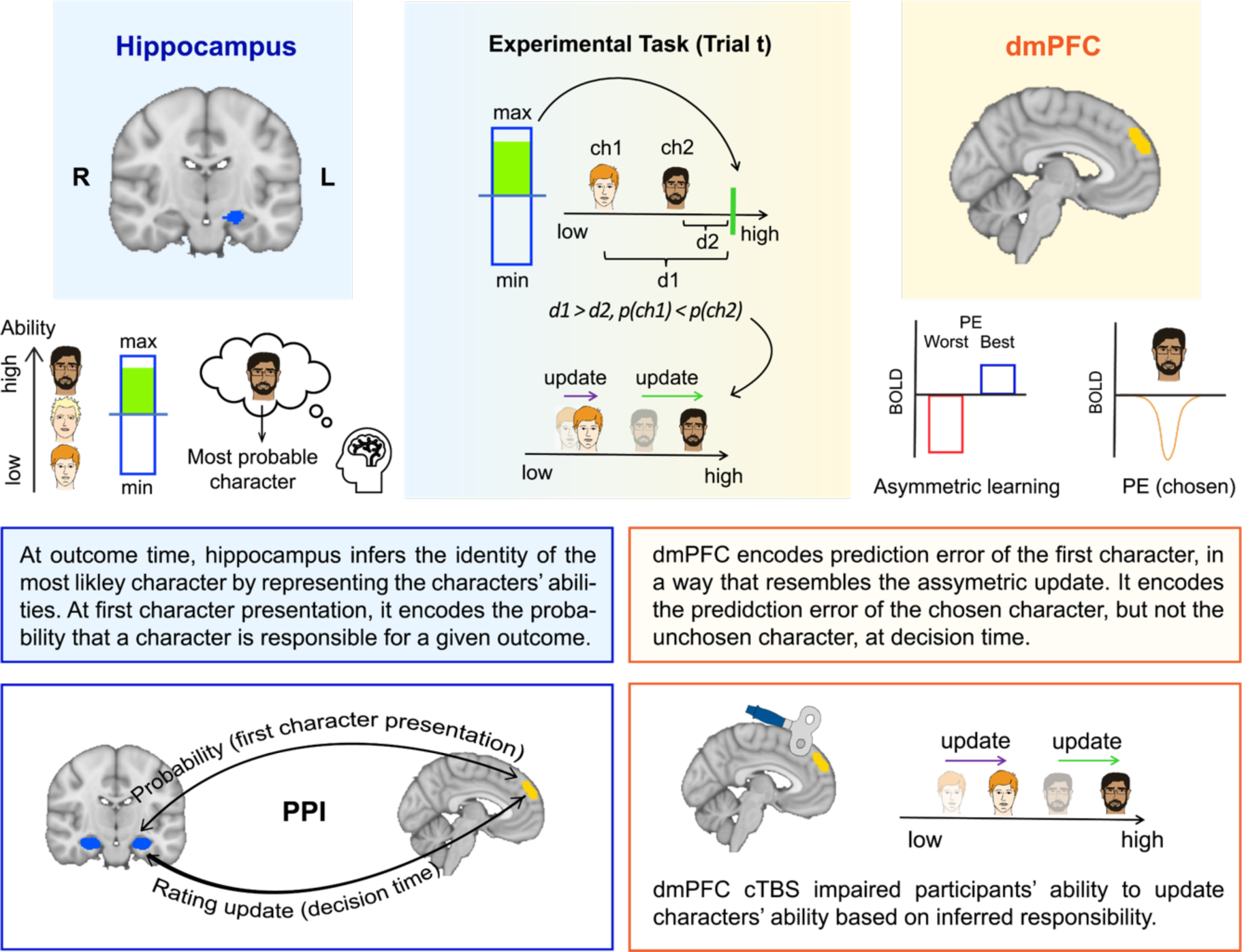
Summary. dmPFC and hippocampus played distinct but complementary roles in latent social cause inference.

Dorsomedial prefrontal cortex (dmPFC) was the second region which played a prominent role in our task. When the first character appeared, its activity was jointly modulated by probability and prediction error (PE): its activity was highest when the best character appeared, and the PE was positive when the worst character appeared, and the PE was negative (Figure 3a). This pattern of activity resembled the asymmetric UEA apparent in behaviour: the update of the better character was larger following positive PEs and the update of the worse character was larger following negative PEs (Figure 2d). This area then encoded the PE of the chosen character (significantly more than the PE of the unchosen character), at the time of choice (Figure 3d). These observations suggest that dmPFC updated the ability estimate for the chosen character (UEA) as a function of the responsibility that participants had inferred the character to have had (LCI). This role of dmPFC is consistent with other studies that have shown that dmPFC learns and represents others’ abilities^15–17^. However, social learning in previous studies was investigated in situations where the cause of each outcome was unambiguous and observable and did not have to be inferred. Our study goes beyond these previous reports and shows that dmPFC is not restricted to updating representations of the ability of others but to doing so in the same asymmetric manner as was evident in behaviour --in a way that was contingent on LCI even though dmPFC did not perform LCI *per se*.

A recent study in humans, using a non-social task found that mPFC represented latent causes in the environment and assigned credit to the inferred cause^40^. Another study found that mPFC inactivation in mice impaired their ability to infer their state and altered dopaminergic reward prediction error signals. The impact was only apparent when the animals’ task state was hidden and needed to be inferred^41^. Our study provides strong evidence for the suggestion that state representations in mPFC and OFC are constructed by hippocampus and relayed to prefrontal areas^3,4,13,26,41^. This claim is supported by several observations: first, the decoding accuracy of the most likely character at the time of outcome, a clear and first possible signature of LCI-related computation, was only observed in hippocampus (Figure 4a). Second, at the next point in the trial, at the first character presentation, the probability that the character was responsible for the outcome was only represented in the hippocampus (Figure 4d). And third, the functional coupling between dmPFC and HPC increased as a function of the probability that the first character was responsible for the outcome (Figure 5a). The hippocampus played this role by representing and tracking participants’ abilities, potentially through its interaction with dmPFC. Indeed, we found that at the time of the second character, when the learning happens, functional coupling between hippocampus and dmPFC increased as the rating update of the chosen character increased (Figure 5b), suggesting hippocampus updates its representation through interaction with dmPFC.

Our computational modelling results indicated that participant behaviour was consistent with a probabilistic model that updates its estimates of characters’ abilities based on the inferred cause of outcome. We modelled this using a combination of two standard Bayesian models from machine learning that have each been extensively used in isolation in neuroscience, Kalman filter and hidden Markov model^2,20–22,42–44^. By leveraging these approaches, the combined HMMKF model computes the probability that each character is accountable for the outcome and adapts its estimates of the responsible character accordingly. However, our preoccupation here is not with the precise details of the HMMKF model but rather with two general features that it possesses. The HMM component discovers the latent cause of each outcome and allows the KF component to update its estimate of the characters on the basis of the inference rather than updating both equally. We argue that any models that are to explain behaviour and neural activity should incorporate these two elements, LCI and UEA, in some way.

Disrupting dmPFC activity could have led to different patterns of impairment. It could have impaired the process of selecting the responsible character while asymmetric UEA could have remained intact. Such a pattern of impairment would have suggested that dmPFC performed LCI and was analogous to the HMM component of our computational model. Alternatively, dmPFC disruption could have preserved participants’ abilities to choose the responsible character, while impairing their ability to update the ability of the responsible character in light of the new outcome. Such a pattern of impairment suggests that dmPFC is KF-analogous and critical for UEA. Our cTBS results confirmed the latter perspective: dmPFC disruption impaired participants’ ability to update character ability estimates as a function of their responsibility. These results suggest that while dmPFC was not necessary for LCI, it played a crucial role in UEA contingent on LCI.

Inferring latent social causes during social interaction entails discovering the latent cause and updating the association between the observation and the cause. Our study provides evidence that the human brain solves and implements this two-stage process through an interaction between hippocampus and prefrontal cortex. These areas play functionally distinct but complementary roles in solving the latent social cause inference problem.

## Methods

### Participants

#### Experiment 1

A total of 37 healthy volunteers without current psychiatric conditions participated in the fMRI study. Four participants were removed due to excessive head motion (i.e., absolute mean displacement > 2.5 mm or relative mean displacement > 0.25 mm), resulting in an effective sample size of 33 participants (mean age±standard deviation=24.55±5.68, 18 female). All participants were right-handed and provided informed consent before the experiment. Participants were paid £40 for their participation in the study, plus up to £35 bonus which was calculated based on their performance.

#### Experiment 4

For experiment 4 (cTBS), 20 participants were recruited. One participant was excluded after the first visit as the required intensity (.8 multiplied by their active motor threshold) was above 50% of the transcranial magnetic stimulation machine’s total output, resulting in a sample size of 19 participants (mean age±standard deviation=20.84±2.29, 10 female). All participants were right-handed and provided informed consent before each session. Participants were paid £100 for their participation in the study, plus up to £30 bonus based on their performance. The study was approved by the Central University Research Ethics Committee (CUREC) of the University of Oxford (Experiment1: R60547/RE001, Experiment2: R76473/RE001).

### Experimental procedure

#### Experiment 1 (fMRI)

Participants underwent a single fMRI session. Participants were required to complete two tasks inside a 3T MRI scanner: the latent social cause inference task and a localiser task (see below). The entire MRI scan, including a structural MRI scan, lasted approximately 80 minutes.

#### Latent social cause inference task

On each trial, an outcome was displayed for 2 seconds using a score frame in the middle of the screen. The top and the bottom of the frame represented the maximum and minimum possible outcome that could have been achieved by the teams, respectively. A horizontal line represented the mid-line (Figure 1a) so outcomes above or below the middle line indicated above- or below-average team performance respectively. In each trial, two of the three characters were pseudorandomly chosen to pair up as a team. After 2 seconds, participants were presented with the first character for a variable amount of time (3.5 to 5.5 seconds) which was generated using a truncated exponential with a mean of 4.5 seconds. Afterwards, the picture of the second character was added to the screen. Participants were allowed to choose the character they thought was responsible for the outcome from the onset of the second character. The task instruction that was given to the participants can be found in the supplementary material (supplementary text 1).

The experiment consisted of 3 blocks of 72 trials each. On each trial, the outcome for each pair of characters (three pairs in total) was generated from three distinct but overlapping distributions with means (48 (for the best and the middle characters), 40 (for the best and the worst characters), and 32 (for the middle and the worst characters) and variance of approximately 100. The outcome associated with each pair was repeated twice in each block on non-consecutive trials. For the first 36 trials in each block, we presented each of the three possible pairs of characters on 12 trials. For the second 36 trials of the block, the same outcome and pairs of characters appeared but the order of presentation of the first and the second characters was reversed. For example, if in trial one outcome *x* appeared and *ch*1 was the first character and *ch*3 was the second character, in trial 37, outcome *x* appeared while *ch*3 was the first character and *ch*1 was the second character. The second block was identical to the first block. In the third block, we exchanged the performance levels of the first and third characters with one another (thereby effecting a reversal in performance level), while keeping all other aspects of the task intact.

In some trials, the choice stage was followed by a rating phase. In the rating phase, participants were required to rate one of the characters on a –5 (min) to +5 (max) scale with 10 steps. Participants rated each character once every nine trials. The rating trials were randomly distributed within these 9 trials. The character whose rating was required was always one of the two characters presented in that trial. The intertrial interval (ITI) was drawn from a truncated exponential distribution between [3.5-5.5] seconds with a mean of 4.5 seconds.

To avoid potential gender bias, we used female characters for female participants and male characters for male participants. The pictures associated with different characters in terms of their abilities were counterbalanced across participants.

#### Localiser task

In the localiser task, participants were required to pay attention to the images that appeared on the screen. On each trial, an image of one of the three characters used in the latent social cause inference task was presented in pseudorandom and interleaved order for 2 seconds, followed by a jittered ITI (drawn from a truncated exponential distribution between 3.5-5.5 seconds with a mean of 4.5 seconds). Each character was presented for 30 trials resulting in 90 trials in total. In some trials, after the presentation of an image, participants were presented with the image of the same character and another character and were asked to identify which character they had just seen. This procedure was included to ensure that the participants paid attention to the images.

### Experiment 4 (TMS)

The TMS experiment included five sessions, a “taster” session (Session 1), a structural MRI session (Session 2), and three experimental sessions (Session 3 to 5). In two of the three experimental sessions, participants received active continuous theta burst stimulation (cTBS) immediately prior to completing the latent social cause inference task, while in the other experimental session, no stimulation was applied (sham session).

**In Session 1**, following face-to-face TMS safety screening, we assessed participants’ active motor thresholds to determine the appropriate intensity of cTBS stimulation for subsequent sessions. Participants whose 80% of active motor thresholds exceeded 50% of the transcranial magnetic stimulation machine’s maximum output were excluded from subsequent sessions (n=1). Additionally, in Session 1, we applied a 10-second “taster” of cTBS over the approximate location of dmPFC with the stimulator output set to 30% of the machine output to familiarize participants with the procedure for the subsequent cTBS sessions.

**In Session 2**, a 3T structural MRI scan was obtained for neuronavigation.

**The remaining three sessions** were experimental task sessions (Session 3, 4, and 5) during which participants performed the latent social cause inference task. In the sham sessions, no stimulation was applied, while in the active stimulation sessions, cTBS stimulation over dorsomedial prefrontal cortex (dmPFC) or a control area (vertex) was applied before starting the latent social cause inference task (see “cTBS protocols” for more details). The order of the three sessions was counterbalanced across participants. The order of the trials (i.e., outcomes and characters that they observed) for all three sessions was identical. However, to avoid any influence of previous sessions, we used different characters for each session. Therefore, in each session, participants learnt the performance of three novel characters. In addition, we changed two other aspects of the latent social cause inference task for the cTBS experiment. First, we made the task easier by making the means of each character’s performance distribution further apart from each other (M=12, 21, 30) while the standard deviations associated with each character’s performance remained unchanged (SD=7). Additionally, we removed the jitters that were introduced for the fMRI experiment. Specifically, the outcome bar was presented for 1 second, followed by the first character presentation for 1.5 seconds. The ITI was set to 2 seconds. The tasks were programmed and presented using the Psychophysics Toolbox in MATLAB^45^.

### cTBS protocol

We used a Magstim-rapid-2 stimulator (MagStim, Whitland, Carmarthenshire, UK) connected to a 70mm figure-8 coil for all transcranial magnetic stimulation (TMS) procedures. TMS was applied in the taster session (Session 1) when assessing participants’ motor active threshold and in the experimental sessions (Session 3-5) in Experiment 4.

During Session 1, we measured the active motor threshold (AMT) of the participants for the left primary motor cortex (M1) “hotspot”. The “hotspot” refers to the specific area on the scalp where TMS elicits the largest motor-evoked potential (MEP). The AMT was defined as the minimum stimulation intensity required to generate an MEP in the right first dorsal interosseous (FDI)^46^, in at least 50% of the trials, while participants exerted constant pressure (10% of their maximum force) between their index finger and thumb. Bipolar surface Ag-AgCl electrodes were used to record the electromyographic (EMG) activity in the right FDI. The recorded responses underwent bandpass filtering between 10 and 1000Hz, with an additional 50 Hz notch filtering. The data were sampled at a rate of 5000 Hz and captured using a D440 Isolated EMG amplifier from Digitimer, a Hum Bug 50/60 Hz Noise Eliminator from Quest Scientific, a CEDmicro1401 Mk.II A/D converter, and a PC running Spike2 software from Cambridge Electronic Design.

During each experimental session (Session 3-5), we applied cTBS to either dmPFC or a control area (vertex) or sham (where no stimulation was applied). We aimed to disrupt activity in the dmPFC which was identified by our fMRI study (Experiment 1; see Figure 3a; MNI x/y/z coordinates in mm: −2/48/38). The vertex was defined as the intersection of the central sulci from both cortical hemishpheres^47^ (MNI /x/y/z coordinates in mm: 0/-34/72), which served as a control stimulation site. Both stimulation sites were first defined in standard MNI space and were then warped to individual structural space using FMRIB Software Library’s (FSL)^48^ non-linear transformations (FNIRT).

A standard neuronavigated cTBS protocol^36^ was applied during each experimental session (except for the sham session where no stimulation was delivered). The stimulation site was projected onto the individual high-resolution, T1-weighted structural scan (Brainsight; Rogue Research). The nasion, nose tip, left ear, and right ear were used for registration of the structural image. During stimulation, the TMS coil was held in place tangentially to the skull by an experimenter. The standard cTBS stimulation protocol comprised 600 pulses in bursts of three pulses at 50 Hz that were applied every 200 ms (total stimulation duration was approximately 40 s). The stimulation intensity was determined by taking 80% of the output of the TMS machine at the participant’s AMT. The use of such a relatively low subthreshold intensity offered the benefit of minimizing the spread of stimulation beyond the intended target area. Participants were instructed to start the task right after the stimulation and minimise their movement while performing the task.

### Behavioural data analysis

#### Multilevel regression models

In all multilevel models, we considered participants as the random variable. To fit the multilevel models, we followed the guidelines proposed by Bates and colleagues^49^ as follows: we started from the full model (the model which contained all possible random effects) and gradually removed the random effect variables until the model converged. A principal component analysis (PCA) was performed on the reduced model to check the dimensionality of the variance-covariance matrix, and a model comparison was conducted in order to assess whether removing certain components would reduce the goodness of fit. We could then decide whether or how to further simplify the random-effects structure based on these two results. The above steps were repeated until an optimal model was identified. All models were performed using lme4 (version 1.1.27.1) package in R^49^. The final models for each analysis were as follows:

##### LM1

This model was conducted to investigate the effect of the difference in characters’ unsigned prediction error (|PE|) on their choice. With this aim, the model was formulated as follows:

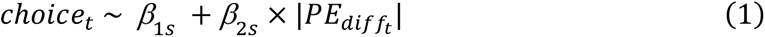

where |*PE_difft_*| is the difference in the unsigned |PE| between the first and the second characters on trial *t*, and *choice_t_* indicates participants’ choice on trial *t* and was set to 1 if the first character was chosen as the responsible character and 0 otherwise. The intercept (*β*_1*s*_) and all slopes with the “s” subscript (e.g., *β_ks_*) were allowed to vary across participants by including random effects of the form *β_ks_* = *β*_*k0*_ + *b_ks_* where 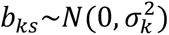.

##### LM2

This model was conducted to investigate the effect of outcome, unsigned prediction error (|PE|) and their interaction on participants’ rating of a character on each trial. With this aim, the model was formulated as follows:

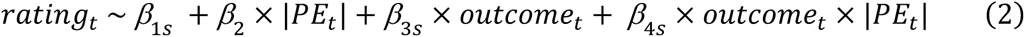

Where *rating_t_* indicates participants’ rating of a character on trial t. |*PE_t_*| and *outcome_t_* indicates the unsigned prediction error and outcome on trial *t*, respectively. The intercept (*β*_1*s*_) and all slopes with the “s” subscript (e.g., *β_ks_*) were allowed to vary across participants by including random effects of the form *β_ks_* = *β*_*k0*_ + *b_ks_* where 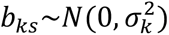.

##### LM3

This model was conducted to investigate the effect of responsibility, the signed PE, and their interaction on participants’ update of a characters’ rating on each trial. With this aim, the model was formulated as follows:

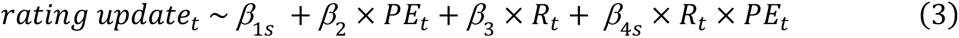

Where *rating update_t_* indicates participants’ update of a character’s rating minus the last time they rated the current character’s rating. *PE_t_* and *R_t_* indicate prediction error and responsibility on trial t, respectively. *R_t_* was set to 1 if the character whose rating was required was also chosen as the responsible character on that trial and was set to 0 otherwise. The intercept (*β*_1*s*_) and all slopes with the “s” subscript (e.g., *β_ks_*) were allowed to vary across participants by including random effects of the form *β_ks_* = *β*_*k0*_ + *b_ks_* where 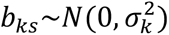.

### TMS data analysis

To test whether dmPFC stimulation might alter participants’ behavioural pattern in the task, we first ran the multilevel models with the same fixed effects structure as the ones employed in Experiment 1 for each condition (i.e., sham, vertex and dmPFC conditions) separately (see Table S6 for details on the model specifications). To compare between different stimulation conditions, we ran each model with only fixed effects for each participant separately and performed paired samples Wilcoxon tests to investigate whether the effects of interest were different between conditions.

We also tested whether the rating difference between the best and worst characters was different across conditions. We used identical task sequences in all three sessions (each character was rated once in each unit comprised of nine trials). This allowed us to calculate the rating difference between the best and the worst characters within each unit. We then compared the rating difference between conditions. The following multilevel model was performed:

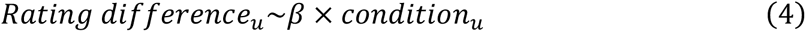

where *Rating difference_t_* denotes the difference in rating between the best and worst characters in unit *u*, and *condition*_u_ is a categorical variable indicating the experimental conditions in unit *u* and was coded as 1 or 2 for vertex and dmPFC, respectively.

### Bayesian statistical analysis

One challenge of interpreting any null result in an fMRI experiment is that it might be due to a lack of statistical power in the fMRI data set (an absence of evidence) rather than the absence of an effect (evidence of absence). To distinguish between these possibilities aim, we employed Bayesian statistics ^50,51^. Bayes Factors are an established tool to assess whether the lack of a significant result is rather due to no effect (i.e. evidence of absence) or a lack of evidence (i.e. absence of evidence) ^28^. We therefore employed Bayesian statistics to examine whether our null results were due to a lack of statistical power or due to a lack of an effect. We computed Bayes Factors (BF) for our key non-significant results using JASP package^28^. We used the categorisation to interpret our BFs ^52,53^.

### Computational modelling

#### Bayesian Learner

We used Bayesian estimation techniques to investigate how a normative model would solve the latent social cause inference task. The model learns the ability of each character by classifying each outcome to one of the characters and updating its estimate of that character’s ability. The model learns a parameter *μ_k_* for each character *k*, which is the characters’ ability. We assume that *μ_k_* is the only unknown parameter and has a known parametric form:

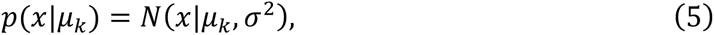

where *x* indicates a given outcome and *σ*^2^ is the variance (constant and known).

We also assume that our prior knowledge can be expressed by a known prior density *p*(*μ_k_*):

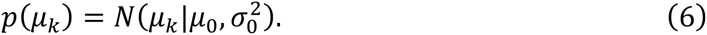

The model then computes the posterior probability given the likelihood and the prior:

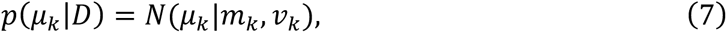

where *m_k_* and *v_k_* are the posterior mean and variance and are calculated as follows:

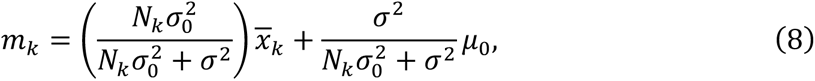

and

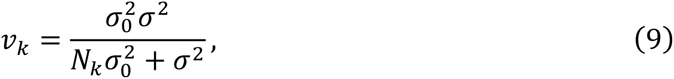

where *N_k_* is the number of trials that the corresponding character was held responsible and *x̄_k_* is the average of outcomes over those trials.

The model tracks *p*(*μ_k_*|*D*) for each character (*k* = 1, 2, 3). After observing an outcome *x* and characters *i* and *j*, the model computes the probabilities *p*(*ch_i_*) = *p*(*μ_i_*|*x*) *and p*(*ch_j_*) = *p*(*μ_j_*|*x*). Using a probabilistic decision rule (a softmax with parameter *β*), the probability of choosing characters *i and j* was proportional to *p*(*ch_i_*) and *p*(*ch_j_*), respectively.

To run simulations for generating Figure 1d and 1e, we ran the model on all sets of trial sequences that we used for participants. Figure 1d presents an illustrative example of the estimates of character’s abilities generated by the model, using one of the trial sequences. When testing the asymmetry in update (Figure 1e), we ran the model on all trial sequences (n=33). As the model chose the responsible character probabilistically, we ran the model 100 times on each trial sequence and computed the mean values across all runs.

#### Kalman Filter

The Kalman Filter (KF) is a well-known Bayesian model for prediction in uncertain environment, which has been widely used in psychology and neuroscience in the past two decades ^20^. The model assumes that outcomes for each character are noisily generated based on a Gaussian generative process that is corrupted by a process noise, *σ* (modelling trial-by-trial noise in the character’s outcome) as well as an observation noise, *ω*. The model updates its previous estimate of the ability of each pair *m*_*t*−1_ following a delta-learning rule after observing the outcome *o_t_* on trial *t*:

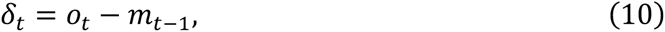

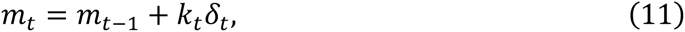

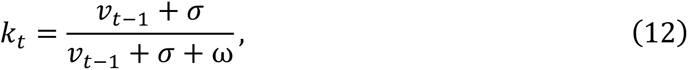

where *k_t_* is the Kalman gain (i.e., learning rate), depending on the posterior variance *v_t_* (i.e., uncertainty) and two noise parameters *σ* and *ω*. The posterior variance *v_t_* is also updated on a trial-by-trial basis:

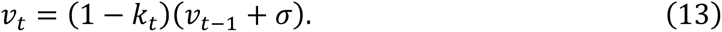

#### Combined Hidden Markov model and Kalman Filter

There was no explicit model of social cause in the Kalman filter approach explained above. This is statistically sound but might not reflect humans’ assumptions in this task. In particular, the Kalman filter model fails to consider the responsibility of characters in updating their corresponding mean and variance (unlike the Bayesian learner model). Therefore, we developed a new model that attributes responsibility to characters and updates its estimate of their ability accordingly. This comes with a further hierarchical inference about the generated outcome for each pair. The model assumes that only one of the characters is responsible for generating the outcome on every trial and update its estimate of the probability of each character being responsible. Thus, there are three latent binary variables encoding the duel between every two characters: *z*_12_ for characters 1 vs 2; *z*_13_ for characters 1 vs 3; and *z*_23_ for characters 2 vs 3. For example, if *z*_12_ is zero when characters 1 and 2 are presented, it attributes responsibility to character 1. Conversely, if *z*_12_ is one, it assigns responsibility to character 2. Additionally, we assume that each character generates the score based on a generative process of the Kalman filter, incorporating noise parameters *σ* and *ω*. Lastly, the generative model assumes that each latent variable *z* may change with a small probability over time depending on parameter *μ*. For example, if *μ* = 0.95, there is 0.95 probability that *z* remains the same as the previous trial, but there is also a small probability (0.05) of it switching. Therefore, the generative process combines the generative process of the Hidden Markov model and the Kalman filter (HMMKF).

For Bayesian inference in this model, we combined particle filtering with Kalman filtering using a well-known method in Bayesian statistics called Rao-Blackwellized particle filtering^42^. Inference over the responsibility parameter is based on the particle filter. The particle filter draws many particles (i.e., samples) of responsibility for the corresponding pair, say *z*_12_, per trial. Given a responsibility sample, the Kalman filter then gives the optimal inference for obtaining the mean and variance for each character. Therefore, for each particle, the model only updates the mean of the responsible character. The particle filter then updates the weight of each particle based on its ability to predict scores. We computed the responsibility assigned to each character by taking a sample from the corresponding responsibility particles (e.g., *z*_12_ if characters 1 and 2 are presented) according to the distribution given by the particle weights. The likelihood was computed using the particle sample and a decision noise parameter, *β*.

### Model fitting and comparisons

Due to the reversal in performance levels between the best and worst performing characters in the last block of the three-block task, we predicted that the model parameters might be different between the first two blocks and the last block. Thus, each model was fitted separately to the first two blocks and to the last block. For fitting the model to the first two blocks, the initial mean *m* and variance *v* for each character’s performance were set to 0 and 1, respectively. For the HMMKF model, 1000 particles were initialised randomly as either 0 or 1, and the weight of each particle was set to 1/1000. For both KF and HMMKF models, the mean *m*, variance *v*, and Kalman gain *k* were updated on a trial-by-trial basis. Therefore, after fitting the model to the first two blocks, we obtained the latest value of the mean *m*, variance *v*, and Kalman gain *k* for each character as well as the updated particles for the HMMKF model. We used the values (*m*, *v*, and *k*) on the last trial of the second block as the initial values of the model for the last block.

Models were fitted to choice data using a maximum *a posteriori* probability (MAP) procedure using the CBM toolbox^54^. The KF and Bayesian model each had one free parameter *β* which was set to be *β* = *e^x^* to ensure positive values for *β*. The HMMKF had one additional free parameter *μ*. For all models, the prior mean and variance of the parameter *x* were set to 0 and 6.25, respectively, similar to the previous studies using this approach^20,54^. For the HMMKF model, *μ* was bounded between 0.5 and 1. We chose the free parameter that best fitted to participants’ trial-by-trial choice quantified by negative log-likelihood.

For model comparison, we used the Variational Bayesian Analysis (VBA) toolbox^23^ in MATLAB R2021b to compare the models using a random-effects approach. The exceedance probability, which measures the probability of a particular model being more likely than all other models^55^ was calculated given the log-model evidence of the models. We also calculated the protected exceedance probabilities, which correct the exceedance probabilities for the possibility that the observed differences in model evidences across subjects might be due to chance rather than meaningful variations^24^.

### fMRI

#### MRI data acquisition

MRI data were acquired using a Siemens 3-Tesla Magnetom Prisma scanner with a 32-channel head coil. Slices were acquired in interleaved order with an oblique angle of −30° from anterior to posterior commissure (AC-PC) to reduce frontal signal dropout. BOLD functional images were collected using multiband gradient-echo EPI sequence with the following parameters: echo time (TE) = 30 ms; repetition time (TR) = 1200 ms; flip angle = 60°; multiband acceleration factor = 3; in-plane (iPAT) acceleration factor = 2; FOV =216mm; voxel size = 2.4 × 2.4 × 2.4 mm. Fieldmap images were acquired to reduce geometric distortion using a dual-echo gradient echo sequence with the following parameters: TE1/TE2 = 4.92/7.38 ms; TR = 482 ms; flip angle = 46°; FOV = 216mm; voxel size = 2.0 × 2.0 × 2.0 mm. Finally, T1-weighted structural images were acquired using an MPRAGE sequence with the following parameters: TE = 3.97 ms; TR = 1900 ms; Inversion time (TI) = 904 ms; 192 sagittal slices; FOV = 192 mm; voxel size = 1.0 × 1.0× 1.0 mm.

#### Preprocessing

Preprocessing of fMRI data was performed using FEAT, part of FSL version 6.00 (FMRIB’s Software Library^56^). We preprocessed the data through the following steps: geometric distortion correction using B0 field mapping (effective echo spacing = 0.275 ms), motion correction using MCFLIRT (FSL)^48^ and slice-timing correction for the interleaved sequence, non-brain extraction using BET^57^, spatial smoothing using a Gaussian filter kernel of FWHM 5.0 mm, and high-pass temporal filtering with 3dB cut-off of 100 s. The functional multiband scans were registered to standard MNI space following a three-step process: (1) linear affine registration (6 degrees of freedom) from the participants’ whole-brain function EPI images to high-contrast functional images (single-band reference) using FLIRT^48^, (2) boundary-based registration from the single-band reference images to the individual high-resolution structural images (T1-weighted) with fieldmap distortion correction; (3) non-linear registration (12 degrees of freedom) from the participants’ T1 structural image to 2 mm standard MNI space using FNIRT^58^.

#### Whole-brain analysis

We performed a three-level whole-brain statistical analyses using FSL’s FEAT^59^. Before analysis, the data were pre-whitened using FILM to account for temporal autocorrelations^60^. At the first level, we applied a univariate general linear model (GLM) approach to each block separately for each participant. All whole-brain GLMs included all phases of a trial as regressors, as follows: the outcome period (a 0.5 s boxcar, from the onset of the outcome); first character (a 2 s boxcar, from the onset of the presentation of the first character); second character/decision phase (a 2 s boxcar, from the onset of the presentation of the second character); choice (a narrow boxcar from the onset of button-press with a duration of 0.1 s); rating phase (a 0.5 s boxcar, from the onset of the rating). All event amplitudes were set to one.

##### GLM1

We performed the first GLM (GLM1) to identify voxels with activity correlated with the prediction error (PE) of the first character. The trial-by-trial estimates of prediction error were included in the model as a parametric regressor, which was time-locked to the onset of the first character presentation.

##### GLM2

The second GLM (GLM2) was designed to test for voxels with activity correlated with the PE of the chosen and unchosen characters. With this aim, two parametric regressors, PE of the chosen character (*PE_chosen_*) and PE of the unchosen character (*PE_unchosen_*) were included in the model, both time-locked to the decision time.

##### GLM3

We performed an additional GLM (GLM3) using the update of the ability (expressed as the probability of being responsible for the outcome) of the first character as the parametric regressor, time-locked to the decision time. This update refers to the difference between the probability estimate for the first character before and after the second character was presented (*p_rev_* − *p_init_*). *p_init_* is the initial probability estimate that a character was responsible for the outcome. For a given character *i* it was calculated as follows:

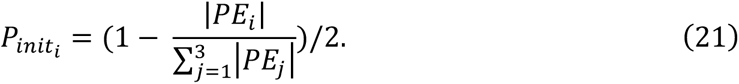

*p_rev_* is the revised probability of the first character being responsible for the outcome at the decision time (i.e., after the presentation of the second character *k*), which was calculated as follows:

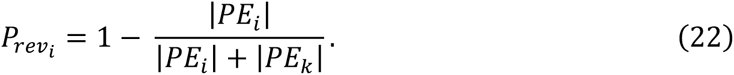

In all GLMS, we denoised the BOLD signal by including the following parameters as additional regressors of no interest: (1) six motion parameters derived from MCFLIRT realignment; (2) regressors removing timepoints with large head motions exceeding a threshold of 0.5 mm for framewise displacement (FD)^61^. Additionally, the temporal derivatives were added for each regressor.

For the second level analysis, we performed within-subject analyses across the three experimental blocks. We averaged the contrast estimates for each participant across three blocks using a fixed-effects approach^59^. Finally, group-level analyses were performed using a mixed-effects approach (FLAME 1+2)^62^ to estimate the average response across participants while accounting for the variance across participants. All results shown in the main effects images were obtained after whole-brain cluster correction with a voxel inclusion threshold of |*Z*|=2.8 and a cluster significance threshold of *p*=.05.

#### Activity time-course analysis

For each time-course analysis, the following steps were first performed to separate the time-course data into epochs: (1) the filtered BOLD time-courses were extracted from each voxel within the ROI/cluster and then averaged across voxels; (2) the nuisance regressors including all the motion parameters were regressed out from the data to reduce the noise in the BOLD signal; (3) the time-course data were normalised and up-sampled by a factor of 10; (4) the up-sampled data were interpolated using the polynomial form of the cubic spline method, and were then epoched in 12 s windows and aligned to the onset that the regressors were time-locked to. Regressors were then fitted to the epoched data at each time step using the ordinary least squares method.

##### TC1.1

This model was used to investigate the effect of PE on dmPFC, vmPFC, and AI/OFC BOLD. Thus, the model was as follows:

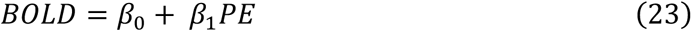

This model was separately run for the best and the worst characters and compared these beta values against each other.

##### TC1.2

This model was used to investigate the effect of the initial probability (*p_init_*) that a character was responsible for the outcome, PE, and their interaction on dmPFC, vmPFC, AI/OFC and hippocampus (HPC).

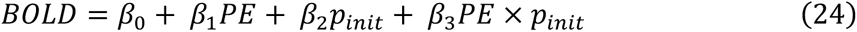

Where *P_init_* is the initial probability that a character was responsible for the outcome. For a given character *i* it was calculated using Equation (21):

Again, this model was separately run for the best and the worst characters and the resulting beta weights were compared against zero.

##### TC1.3

This model was conducted to test whether dmPFC encoded probability of the chosen character at the time of choice. To avoid selection bias, in addition to our dmPFC ROI, we used an anatomical ROI from an independent study encompassing area 9. Thus, the model is as follows:

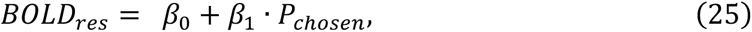

where *BOLD_res_* is obtained by extracting the dmPFC activity time courses from the residuals of GLM2, aligned to the decision time. We took the residuals to control for the colinearity between the PE of the chosen character and probability of the chosen character (*P_chosen_*). *P_chosen_* was calculated as follows:

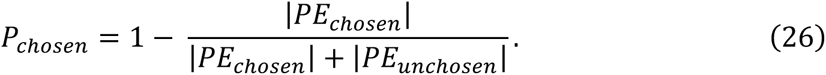

#### Psychophysiological (PPI) analysis

To test functional connectivity between dmPFC and hippocampus, we performed psychophysiological interactions (PPI) analyses. First, we aimed to test whether the relationship between dmPFC and HPC was modulated by the probability that the first character was responsible for the outcome (*P_init_*) at first character presentation. We performed the following time-course analysis using the epoched time course aligned to the onset of the first character presentation:

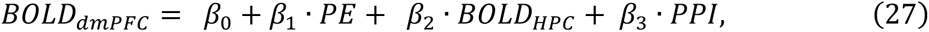

where *BOLD_dmPFC_* is BOLD activity extracted from the a priori ROI over dmPFC (area9), *BOLD_HPC_* is BOLD activity extracted from the left and right hippocampus, and *PPI* is a two-way interaction between *BOLD_HPC_* and PE of the first character.

The second PPI analysis aimed to investigate whether the relationship between dmPFC and hippocampus at decision time was modulated by the rating update for the chosen character. We then performed the following time-course analysis using the epoched time course aligned to the decision time:

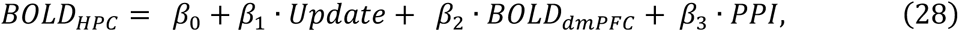

where *PPI* is a two-way interaction between *BOLD_dmPFC_* and rating update for the chosen character.

For all time course analyses, the parameter estimates at each time step were averaged across blocks for every participant using a fixed-effects approach. We tested group-level significance using a leave-one-out procedure to avoid any selection bias. For each participant and for each time course signal, we computed the peak signal (positive or negative) for the group and calculated the beta weight of the left-out participant at the time of the group peak. We repeated this procedure for each participant and compared the resulting beta weights against zero.

### Multivariate Pattern Analysis

#### Decoding Analysis

Both searchlight and ROI-based Multivariate Pattern Analysis (MVPA) analyses were implemented using the pyMVPA toolbox^27^ and custom Python scripts. Prior to the analyses, the raw fMRI data were first motion corrected using FLIRT from FSL^48^, but no spatial or temporal filtering was applied to preserve local voxel information. The data were then detrended to remove linear trend for each scan run of the main task and for the localiser run individually. We used the localiser task as the training set with each character as the label. For this purpose, for each voxel, we first estimated a beta value for each class (character) by fitting a GLM with one regressor time-locked to the presentation of character images. The regressor was defined as a boxcar of 2 seconds in length and convolved with a canonical hemodynamic response function. We then trained the classifier using a linear support vector machine (SVM). We first tested the reliability of the classifier by classifying the identity of the first character at first character presentation. The voxel-wise beta estimate was obtained by fitting an event-related GLM with a regressor at the time of the first character presentation introduced as a boxcar of 3 seconds in length.

After establishing the reliability of the approach, we tested whether the decoding accuracy of the most probable character, following the observation of the outcome, was significantly higher than the chance level in any specific brain region. The probability that a character k (k=1, 2, 3) was responsible for the outcome was calculated as follows:

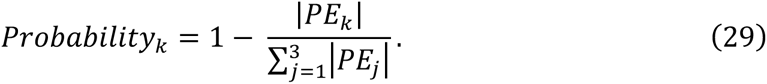

The character with the highest probability was labelled as the most likely character who caused the outcome and was therefore likely to appear next.

The voxel activity patterns tested for this analysis were estimated using another event-related GLM with a regressor at the time of outcome presentation defined as a boxcar of 2 seconds in length. The regressors for both GLMs were convolved with a canonical hemodynamic response function and modelled separately for each scan run. The classification accuracy was measured by taking the proportion of correctly classified trials for each individual run. The classification accuracy was averaged across the three scan runs.

On each trial the classifier returned a class probability for each character. We used the class probability of the most likely character for each trial to compute the decoding accuracy. For the searchlight MVPA, using a whole-brain searchlight method, the above process was repeated for all spheres with a 4-voxel radius, where each sphere was centred around each voxel in the brain once. This searchlight procedure resulted in a decoding accuracy map for each participant. We then performed group-level analyses on these statistical maps to identify regions where the classification accuracy exceeded the chance level (1/3). Specifically, we normalised each participant’s accuracy map from native space to MNI standard space and then performed random-effects t-tests at each searchlight centre using the CoSMoMVPA toolbox^63^ to compare each voxel against the null hypothesis of chance level. The resulting statistical map was voxel-wise thresholded at *p*<.001 and cluster corrected at *p*<.05.

For the ROI-based MVPA, the above analyses were restricted to voxels in each target mask. All the masks were first defined in the MNI space and were then transformed into participants’ native spaces. The resulting decoding accuracy for each ROI were then tested against the chance level using one-tailed Wilcoxon signed-rank tests. The resulting p values were corrected for multiple comparisons using Holm-Bonferroni correction (HBC).

#### Representation Similarity Analysis (RSA)

MRI data were analysed using Matlab (2020b) and Statistical Parametric Mapping software (SPM12; Wellcome Trust Centre for Neuroimaging, London, UK). Preprocessing comprised motion correction, correction for field inhomogeneity, spatial smoothing (5mm FWHM Gaussian kernel), realignment, coregistration, and normalisation to the MNI template and high-pass filtering (128 seconds) following SPM12 standard procedures.

We performed representational similarity analysis (RSA)^32,33^ to test whether the ROIs carried information about characters’ abilities in the task at the time of outcome. To estimate voxel activity patterns, we constructed an event-related GLM with the following regressors: one regressor locked to the outcome screen introduced as a boxcar of 2 seconds in length, one regressor at the time of the first character presentation introduced as a boxcar of 2 seconds in length, one regressor for the presentation of the second character defined as a boxcar of 2 seconds in length, one regressor for button press introduced as an impulse function, one regressor for the presentation of rating (only in trials which rating was required) introduced as a boxcar of 2 seconds, and finally one regressor for button press of rating trials introduced as an impulse function. Regressors were convolved with a canonical hemodynamic response function. Regressors were modelled separately for each scan run, and constants were included to account for differences in mean activation between runs and scanner drifts.

We created a neural representational dissimilarity matrix (RDM) for each scan run. This RDM was constructed using the Mahalanobis distance (Euclidean distance after multivariate noise normalization) between the activity patterns for all 72 trials of each run^64^. Since the neural RDM was constructed using activity patterns from the same scan run (symmetric) and not different scan runs (cross-validated), we only used the upper triangle (and not the lower triangle and diagonal) of the resulting 72 x 72 neural RDM.

To test for the representation of characters’ abilities, we first for each trial obtained the model-derived ability estimate for each character resulting in a vector of abilities for each trial. This vector contained three values (ability estimate for the three characters). We constructed our model RDM by computing the pairwise Euclidian distance between these vectors across trials.

We used rank correlation (Kendall’s *τ_A_*) to quantify the extent to which pattern activity within each ROI was explained by a model RDM. Specifically, we first computed the correlation between the model and neural RDMs for each scan run and then averaged the correlations across the three runs for subsequent group analysis. As described above, we only used the upper triangle of the RDMs for analysis. We used one-tailed Wilcoxon signed-rank tests to test the correlations against 0 and employed Holm-Bonferroni correction (HBC) to account for multiple comparisons.

### Exemplar dissimilarity index (EDI)

The EDI tests whether an ROI carries information about a set of conditions^29^. For this analysis, condition was defined as different characters. To this end, at the time of the first character, we first modelled each character as a separate regressor, separately for each run. We then constructed a split-data Representational Dissimilarity Matrix (sd-RDM) for each scan run (a 3 by 3 matrix as we had three different characters). For each run (three runs in total), each entry of this matrix contained the Euclidian distance between the activity pattern of condition *i* of this run and the average activity pattern of condition *j* of the two remaining runs. We then averaged the three resulting matrices and computed the sum of the diagonal entries (within condition comparisons) minus the sum of the off-diagonal entries (between condition comparisons). For group-level comparison, we used one-tailed Wilcoxon signed-rank tests to test the resulting values against 0.

### Mask for ROI analysis

For dmPFC, we used an anatomical mask (encompassing area 9) based on a previous study which performed parcellation of medial frontal cortex ^25^. We obtained our hippocampus mask from a a previous study^65^. We selected the OFC ROI using the following labels from the automated anatomical labelling atlas 3^66^(AAL3): bilateral medial, anterior, posterior, and lateral orbital gyrus, and bilateral rectal gyrus.

## Supplementary information

**Figure S1.**
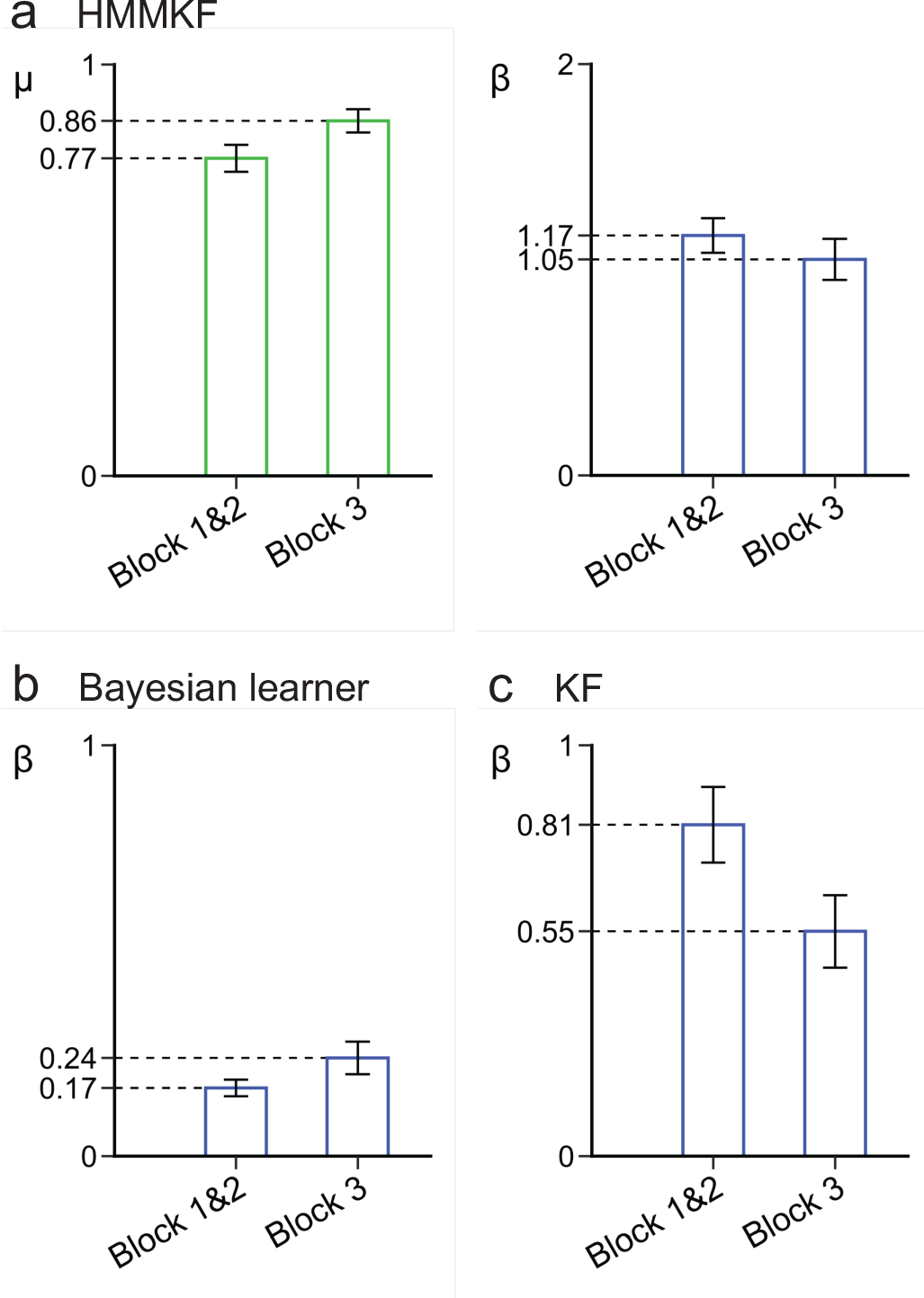
Fitted values for the three computational models. **a** μ and β for the HMMKF. **b** β for the Bayesian learner. **c** β for the KF model. Error bars indicate the standard error of the mean (s.e.m).

**Figure S2.**
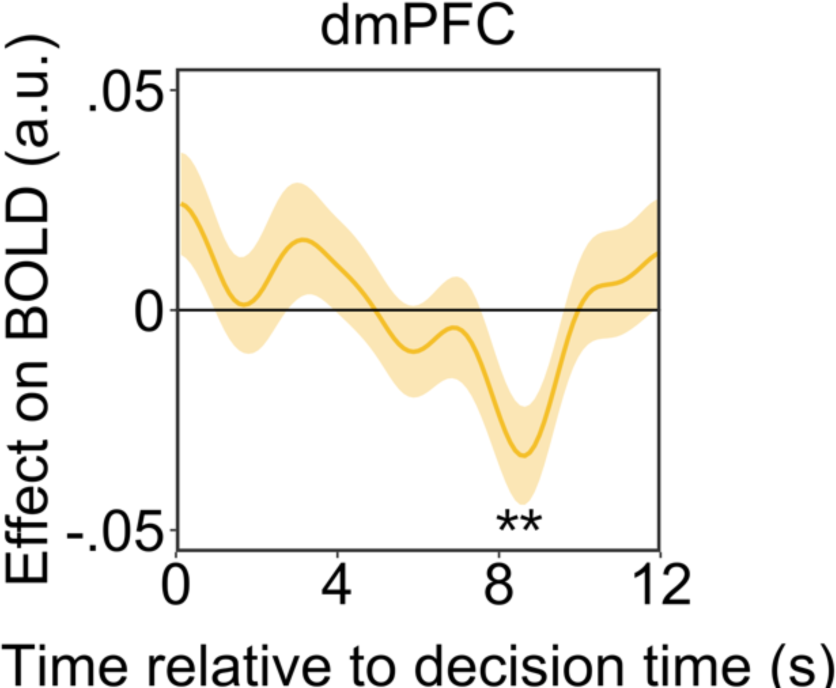
dmPFC encodes the probability that the chosen character is responsible for the outcome. In the main text (Figure 3g), we showed that area 9 encoded *p_chosen_*, here we show that we can replicate this result, if we use the ROI that we found in this study to encode PE of the first character (*W*=128, *p*=.007).

**Figure S3.**
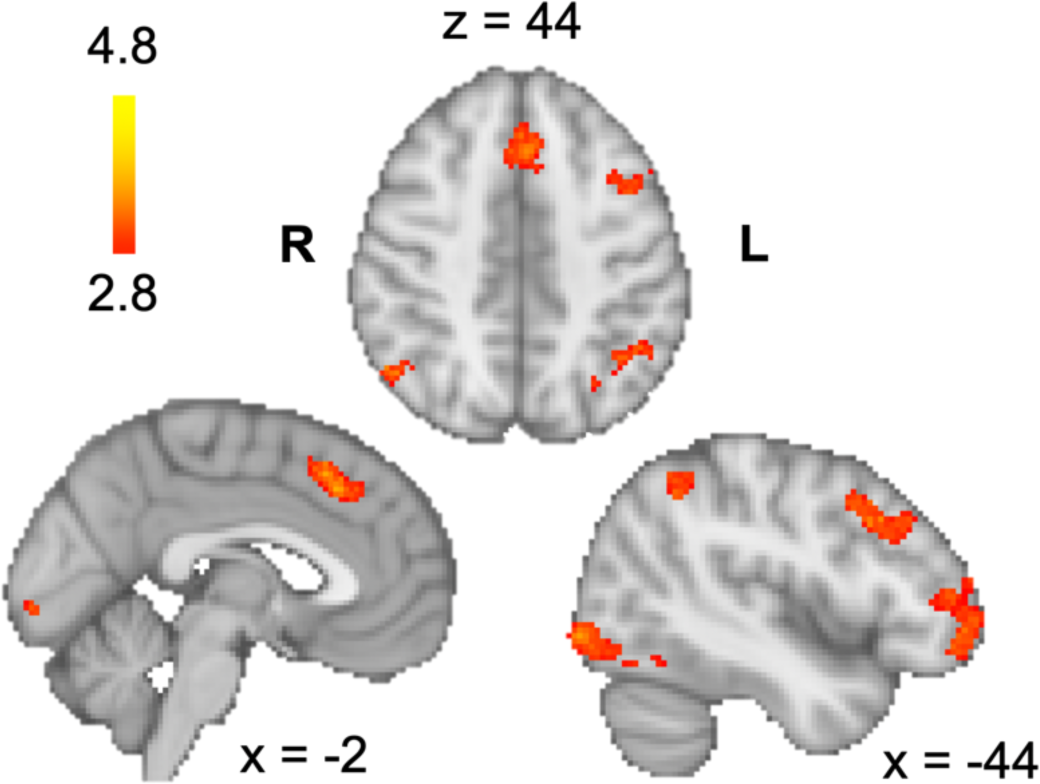
Brain regions signalling decision variable. Whole-brain analysis (Z>2.8, whole-brain cluster-corrected at *p*<.05) revealed brain areas which encoded the decision variable (the difference between the unsigned PE of the chosen and unchosen character) at the decision time, including dorsal lateral prefrontal cortex (dlPFC; peak Z=3.86, MNI: x=-44, y=20, z=36), frontal pole (peak Z=3.85, MNI: x=-46, y=52, z=-6), and pre-supplementary motor area/dorsal anterior cingulate cortex (pre-SMA/dACC; peak Z=4.16, MNI: x=-2, y=24, z=42). All significant clusters are summarised in Table S4.

**Figure S4.**
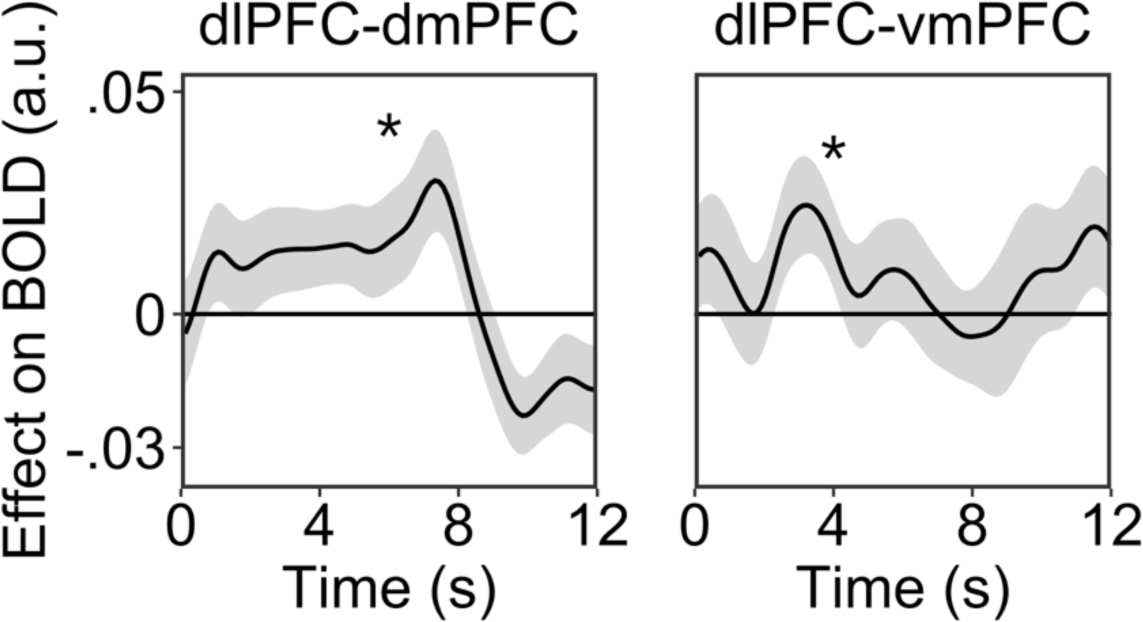
Functional coupling between dlPFC and medial prefrontal cortex (mPFC). PPI analyses showed that at the time of choice, the functional connectivity between dlPFC and dmPFC was modulated by PE of the chosen character (*W*=404, *p*=.014; left panel), and the functional connectivity between dlPFC and vmPFC was modulated by PE of the unchosen character (*W*=379, *p*=.040; right panel).

**Figure S5.**
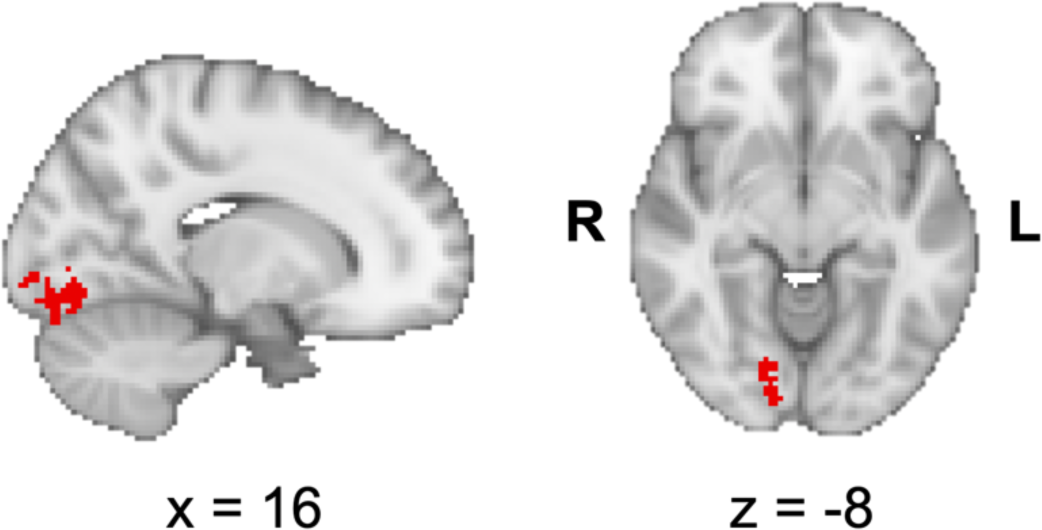
Decoding of characters’ identity in occipital fusiform gyrus at first character presentation. At first character presentation, our whole brain decoding analysis showed that identity of the characters can be significantly decoded from activity pattern of a cluster in the fusiform gyrus (*Z*=2.29; voxel-wise threshold at *p*<.001, cluster correction at *p*<.05). In other words, when the characters are directly perceived, their identities are encoded in extrastriate visual cortex but when their identities are inferred from an outcome for which they might be responsible, their identities are encoded in hippocampus.

**Figure S6.**
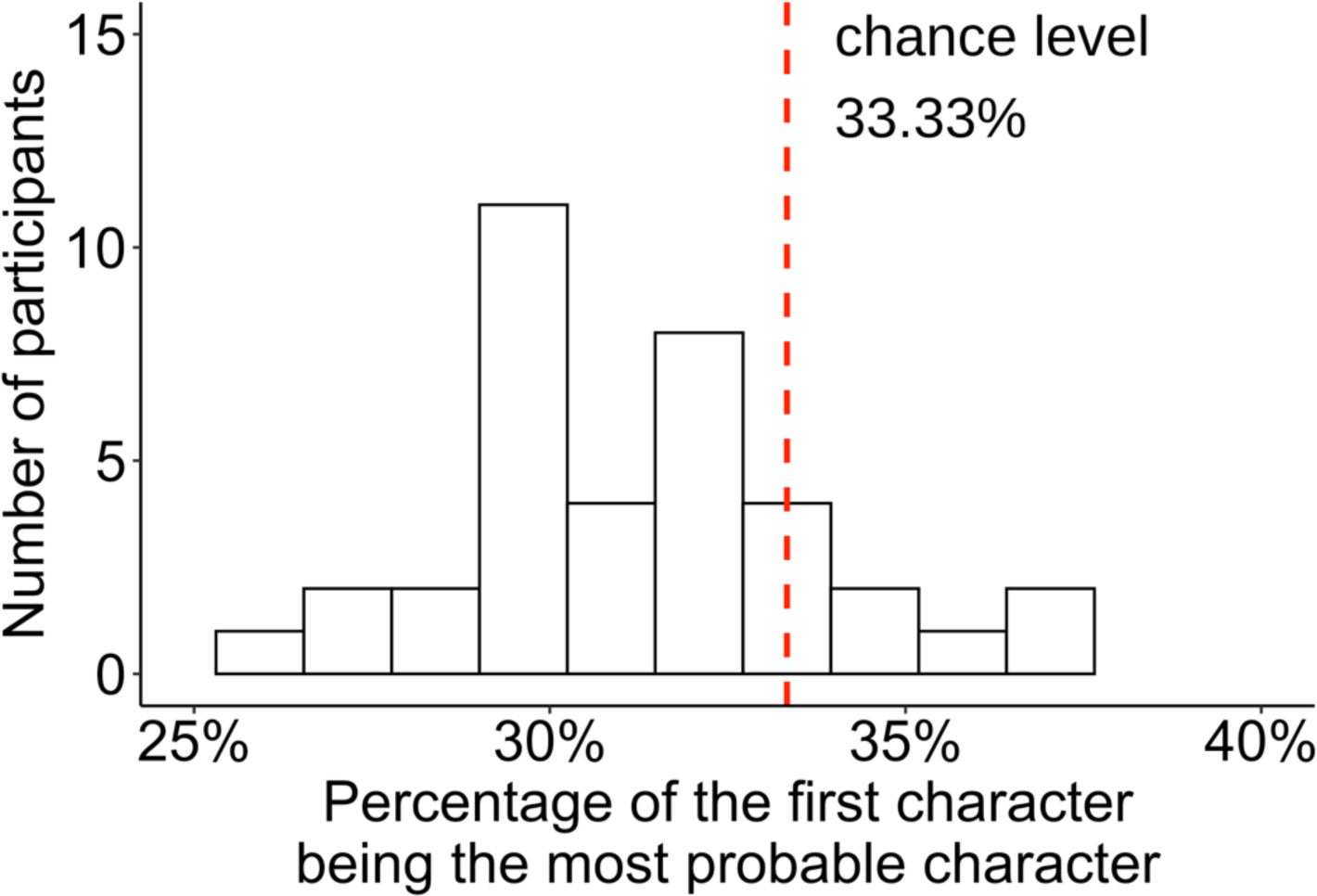
The percentage of occasions on which the first character was the most probable character after observing the outcome was below chance level of 33.33% (*W*=86, *p*<.001), related to Figure 4a.

**Figure S7.**
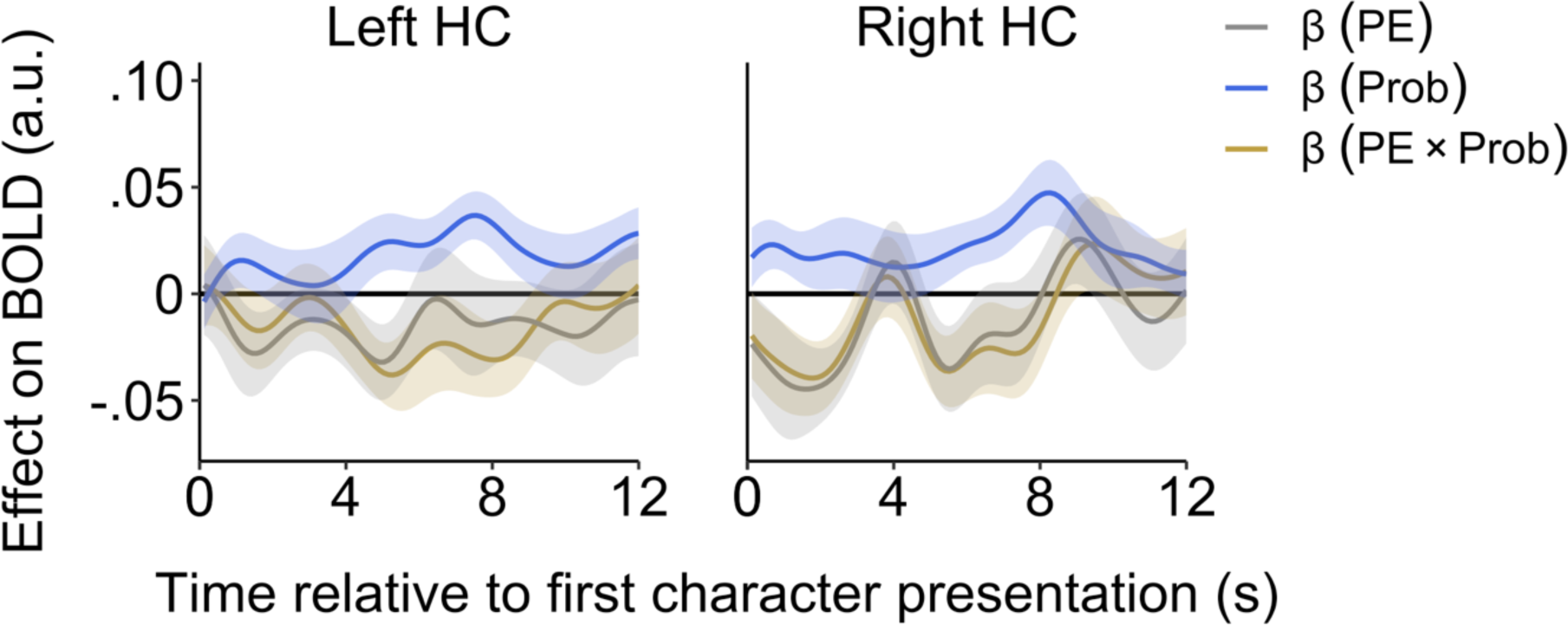
In the main text (Figure 4d), we showed that hippocampus encoded the probability that the first character was responsible for a given outcome (*p_init_*). Here we show that, unlike dmPFC, hippocampal activity was not modulated by the PE of the first character (Left HPC: *W*=190, *p*=.108; Right HPC: *W*=200, *p*=.270; HBC) or the interaction between PE and *p_init_* (Left HPC: *W*=203, *p*=.338; Right HPC: *W*=170, *p*=.123; HBC).

**Figure S8.**
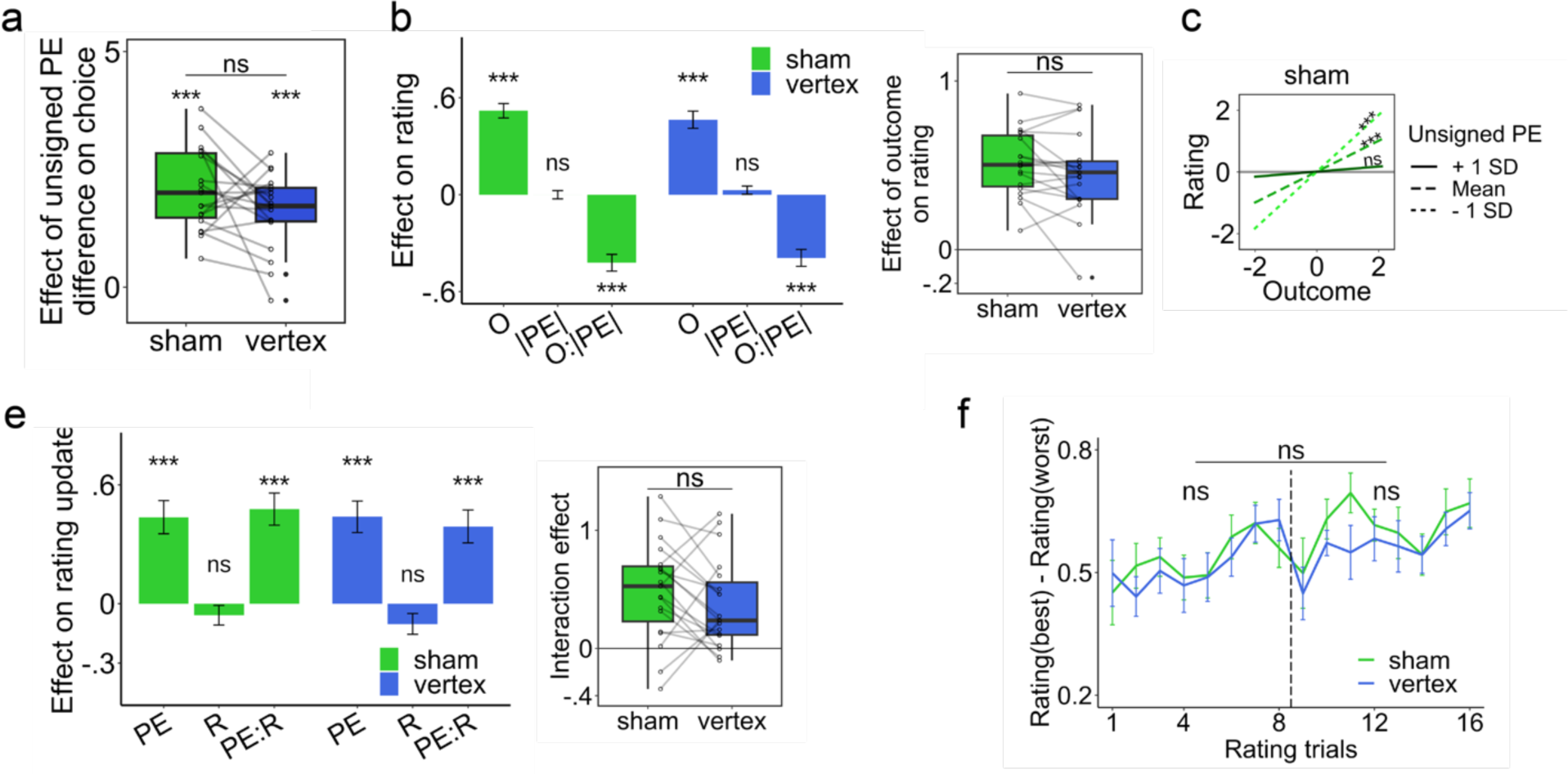
Supplementary cTBS results (sham vs. vertex), related to Figure 6. **(a)** The effect of unsigned PE on choice was quantified using a regression model. This effect was not different between control and vertex conditions; **(b)** Using a multilevel regression model, we quantified the effect of unsigned PE, outcome, and their interaction on rating. The effect of outcome on rating was not different between control and vertex conditions; **(c)** The effect of outcome on rating is plotted for large, medium and small unsigned PEs. Unlike in dmPFC condition (Figure 6d), the effect of outcome on rating is not significant for large PEs in both control and vertex condition; **(d)** Using a multilevel regression model, we quantified the effect of PE, responsibility (R), and their interaction on rating update. The interaction effect was not different between control and vertex conditions; **(e)** Rating difference between the best and the worst characters was not different between vertex and sham conditions (Block 1: *B*=-0.01, *SE*=0.03, *p*=.757; Block 2: *B*=-0.05, *SE*=0.03, *p*=.078; when considering two blocks together: *B*=-0.03, *SE*=0.02, *p*=.145). See detailed cTBS results (including regression coefficients and comparisons between conditions) in Table S6.

**Table S1.**
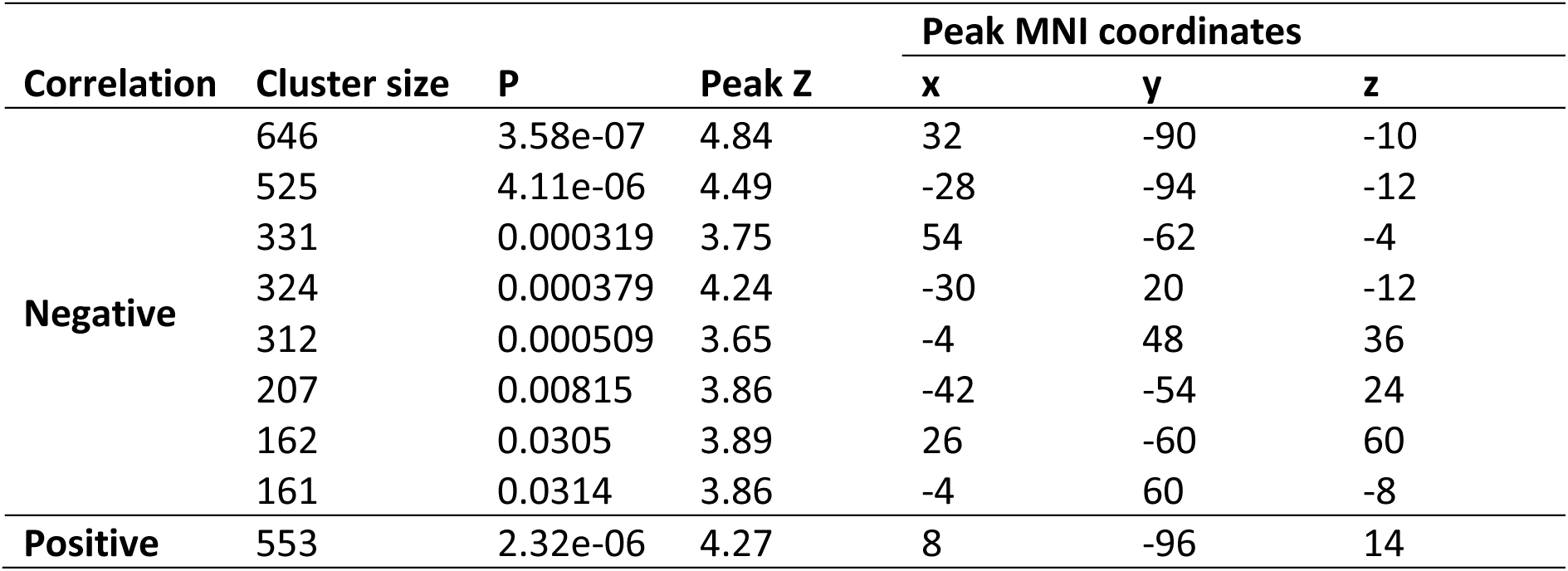
Clusters that significantly correlated with prediction error of the first character at the time of first character presentation, related to Figure 3a.

| Correlation | Cluster size | P | Peak Z | Peak MNI coordinates |  |  |
| --- | --- | --- | --- | --- | --- | --- |
|  |  |  |  | x | y | z |
| Negative | 646 | 3.58e-07 | 4.84 | 32 | -90 | -10 |
|  | 525 | 4.11e-06 | 4.49 | -28 | -94 | -12 |
|  | 331 | 0.000319 | 3.75 | 54 | -62 | -4 |
|  | 324 | 0.000379 | 4.24 | -30 | 20 | -12 |
|  | 312 | 0.000509 | 3.65 | -4 | 48 | 36 |
|  | 207 | 0.00815 | 3.86 | -42 | -54 | 24 |
|  | 162 | 0.0305 | 3.89 | 26 | -60 | 60 |
|  | 161 | 0.0314 | 3.86 | -4 | 60 | -8 |
| Positive | 553 | 2.32e-06 | 4.27 | 8 | -96 | 14 |

**Table S2.** Brain areas that significantly correlated with prediction error of the chosen and unchosen character at the decision time, related to Figure 3d.

| Contrast<br>(Correlation) | Cluster size | P | Peak Z | Peak MNI coordinates |  |  |
| --- | --- | --- | --- | --- | --- | --- |
|  |  |  |  | x | y | z |
| Chosen<br>(Negative) | 3288 | 1.2e-21 | 4.09 | -44 | -20 | 64 |
|  | 1541 | 1.13e-12 | 4.70 | -6 | -74 | 2 |
|  | 1115 | 5.02e-10 | 3.88 | 60 | -20 | 30 |
|  | 951 | 6.42e-09 | 3.88 | 36 | 18 | 10 |
|  | 584 | 3.58e-06 | 3.77 | -30 | -70 | -20 |
|  | 582 | 3.7e-06 | 3.95 | 8 | 18 | 50 |
|  | 383 | 0.000202 | 3.67 | -14 | -64 | 62 |
|  | 362 | 0.00032 | 3.81 | 42 | -84 | 18 |
|  | 341 | 0.000511 | 3.81 | 42 | 40 | 12 |
|  | 327 | 0.000701 | 3.86 | -40 | 34 | 34 |
|  | 274 | 0.00243 | 3.87 | 38 | 0 | 62 |
|  | 179 | 0.028 | 3.63 | 26 | 52 | 32 |
| Unchosen<br>(Negative) | 767 | 2.38e-07 | 3.92 | -6 | -30 | 58 |
|  | 605 | 3.58e-06 | 4.09 | 24 | -84 | -34 |
|  | 285 | 0.00233 | 3.80 | 0 | 40 | -20 |
|  | 232 | 0.0083 | 3.77 | -28 | -96 | -18 |
|  | 225 | 0.00988 | 3.77 | -28 | -84 | -36 |
|  | 186 | 0.027 | 3.57 | -62 | -56 | -4 |
|  | 179 | 0.0325 | 3.90 | -30 | 30 | 50 |
|  | 175 | 0.0361 | 3.78 | -12 | -44 | 32 |

**Table S3.** Brain areas that significantly correlated with the difference between the revised and the initial probability (*p_rev_* − *p_init_*) at the second character presentation, related to Figure 3f.

| Correlation | Cluster size | P | Peak Z | Peak MNI coordinates |  |  |
| --- | --- | --- | --- | --- | --- | --- |
|  |  |  |  | x | y | z |
| Negative | 2897 | 1.06e-28 | 3.93 | -2 | 34 | 54 |
|  | 356 | 3.52e-06 | 3.69 | 50 | -48 | 44 |
|  | 296 | 2.52e-05 | 3.54 | -32 | 60 | 6 |
|  | 260 | 8.76e-05 | 3.80 | 2 | -24 | 28 |
|  | 154 | 0.00511 | 3.64 | -56 | -56 | 38 |
|  | 129 | 0.0151 | 3.58 | -52 | 30 | 10 |

**Table S4.** Brain areas that significantly correlated the decision signal (the difference between the unsigned PE of the chosen and unchosen character) at the decision time, related to Figure S2.

| Correlation | Cluster size | P | Peak Z | Peak MNI coordinates |  |  |
| --- | --- | --- | --- | --- | --- | --- |
|  |  |  |  | x | y | z |
| Positive | 3473 | 1.65e-21 | 4.77 | -30 | -86 | -14 |
|  | 425 | 0.000138 | 3.85 | -46 | 52 | -6 |
|  | 285 | 0.00271 | 4.16 | -2 | 24 | 42 |
|  | 235 | 0.00877 | 3.86 | -44 | 20 | 36 |
|  | 229 | 0.0101 | 4.07 | 46 | -58 | 38 |
|  | 174 | 0.041 | 3.60 | -44 | -54 | 46 |

**Table S5.**
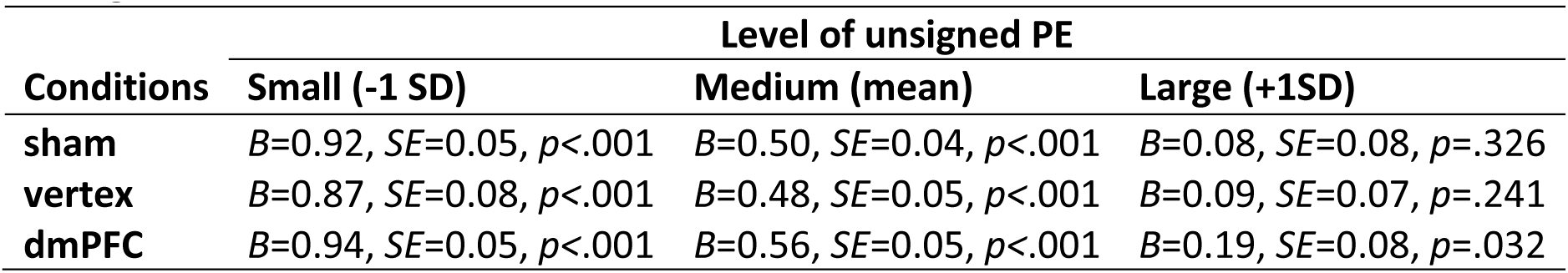
Effect of outcome on rating at different levels of unsigned PE, related to Figure 6d and Figure S5c.

**Table S6.** Detailed cTBS results, related to Figure 6 and Figure S5.

**LM1:** $choice_t \sim \beta_{1s} + \beta_{2s} \times PE\_diff_t$
| Regressor | Coefficient |  |  | Comparison |  |
| --- | --- | --- | --- | --- | --- |
|  | sham | vertex | dmPFC | vertex-dmPFC | sham-vertex |
| **Unsigned PE difference** | $B=1.99,$<br>$SE=0.18,$<br>$p<.001$ | $B=1.59,$<br>$SE=0.18,$<br>$p<.001$ | $B=1.47,$<br>$SE=0.12,$<br>$p<.001$ | $V=116,$<br>$p=.418$ | $V=137,$<br>$p=.096$ |

| Regressor | Coefficient |  |  | Comparison |  |
| --- | --- | --- | --- | --- | --- |
|  | sham | vertex | dmPFC | vertex-dmPFC | sham-vertex |
| **Unsigned PE** | $B=-0.00,$<br>$SE=0.03,$<br>$p=.993$ | $B=0.02,$<br>$SE=0.03,$<br>$p=.959$ | $B=0.05,$<br>$SE=0.03,$<br>$p=.054$ | $V=70,$<br>$p=.332$ | $V=83,$<br>$p=.651$ |
| **Outcome** | $B=0.50,$<br>$SE=0.04,$<br>$p<.001$ | $B=0.48,$<br>$SE=0.05,$<br>$p<.001$ | $B=0.56,$<br>$SE=0.05,$<br>$p<.001$ | $V=23,$<br>$p=.002$ | $V=132,$<br>$p=.145$ |
| **Interaction** | $B=-0.42,$<br>$SE=0.05,$<br>$p<.001$ | $B=-0.39,$<br>$SE=0.05,$<br>$p<.001$ | $B=-0.37,$<br>$SE=0.05,$<br>$p<.001$ | $V=91,$<br>$p=.891$ | $V=84,$<br>$p=.679$ |

| Regressor | Coefficient |  |  | Comparison |  |
| --- | --- | --- | --- | --- | --- |
|  | sham | vertex | dmPFC | vertex-dmPFC | sham-vertex |
| **PE** | $B=0.44,$<br>$SE=0.08,$<br>$p<.001$ | $B=0.44,$<br>$SE=0.08,$<br>$p<.001$ | $B=0.58,$<br>$SE=0.10,$<br>$p<.001$ | $V=49,$<br>$p=.066$ | $V=97,$<br>$p=.953$ |
| **Responsibility** | $B=-0.04,$<br>$SE=0.06,$<br>$p=.460$ | $B=-0.12,$<br>$SE=0.07,$<br>$p=.106$ | $B=-0.05,$<br>$SE=0.06,$<br>$p=.402$ | $V=82,$<br>$p=.623$ | $V=101,$<br>$p=.829$ |
| **Interaction** | $B=0.48,$<br>$SE=0.09,$<br>$p<.001$ | $B=0.38,$<br>$SE=0.08,$<br>$p<.001$ | $B=0.12,$<br>$SE=0.09,$<br>$p=.189$ | $V=148,$<br>$p=.032$ | $V=118,$<br>$p=.374$ |

### Supplementary text 1: Task instruction

**Important note:** Actual human faces that were used for the study were replaced by cartoons here.

**START OF THE INSTRUCTION AS IT APPEARED:**

**----------------**

**INSTRUCTIONS FOR RESPONSIBILITY GAME**

#### Overview of the study

Three individuals previously were paired together in groups of two and completed a minitask for several rounds. The minitask required them to work together as a team. Each time, the two individual received feedback about their performance. In this experiment, you will see each team’s performance for several rounds. <u>The individuals’ contribution to the minitask was</u> <u>different. Some were more competent than others. Their competence also varied from round</u> <u>to round.</u> Even though you cannot see each individual’s contribution to their teamwork, your task is to learn their ability in the task.

Therefore, your task is:

- First, indicate which individual is more responsible for the outcome of their teamwork.
- Second, indicate your estimate of each individual’s performance in the task from time to time.

Each individual is assigned a photo from a public face database in order to keep their identity anonymous.

#### Task

There will be three blocks in total. In each block, there will be 72 rounds (see the structure of the task in Figure 1).

**Figure 4.**
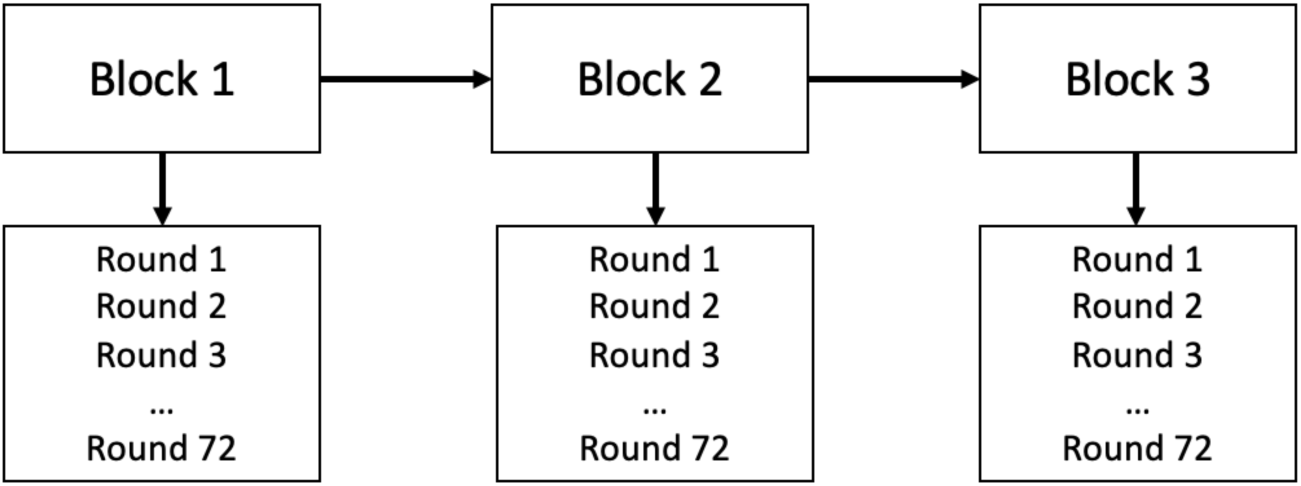

In each round, you will first observe the outcome in the middle of the screen which was the result of the individuals’ teamwork. A vertical score frame is devised to represented the outcome of the individuals’ teamwork (Figure 2). The bottom end of the score frame indicates the minimum score that the teams could have achieved (which is marked by min). The top end of the score frame indicates the maximum score that they could have achieved (which is marked by max). A horizontal line in the middle of the frame indicates its midpoint. A green bar inside the frame indicates the score that a team has achieved.

As you can see in Figure 2, the score frame on the left side shows an amazing outcome because the green bar is near the maximum point. The right one shows a very poor outcome because the green bar is near the bottom of the score frame. Sometimes the outcomes are neither great nor terrible. For example, the one in the middle shows a mediocre outcome because the green bar is near the midpoint.

**Figure 5.**
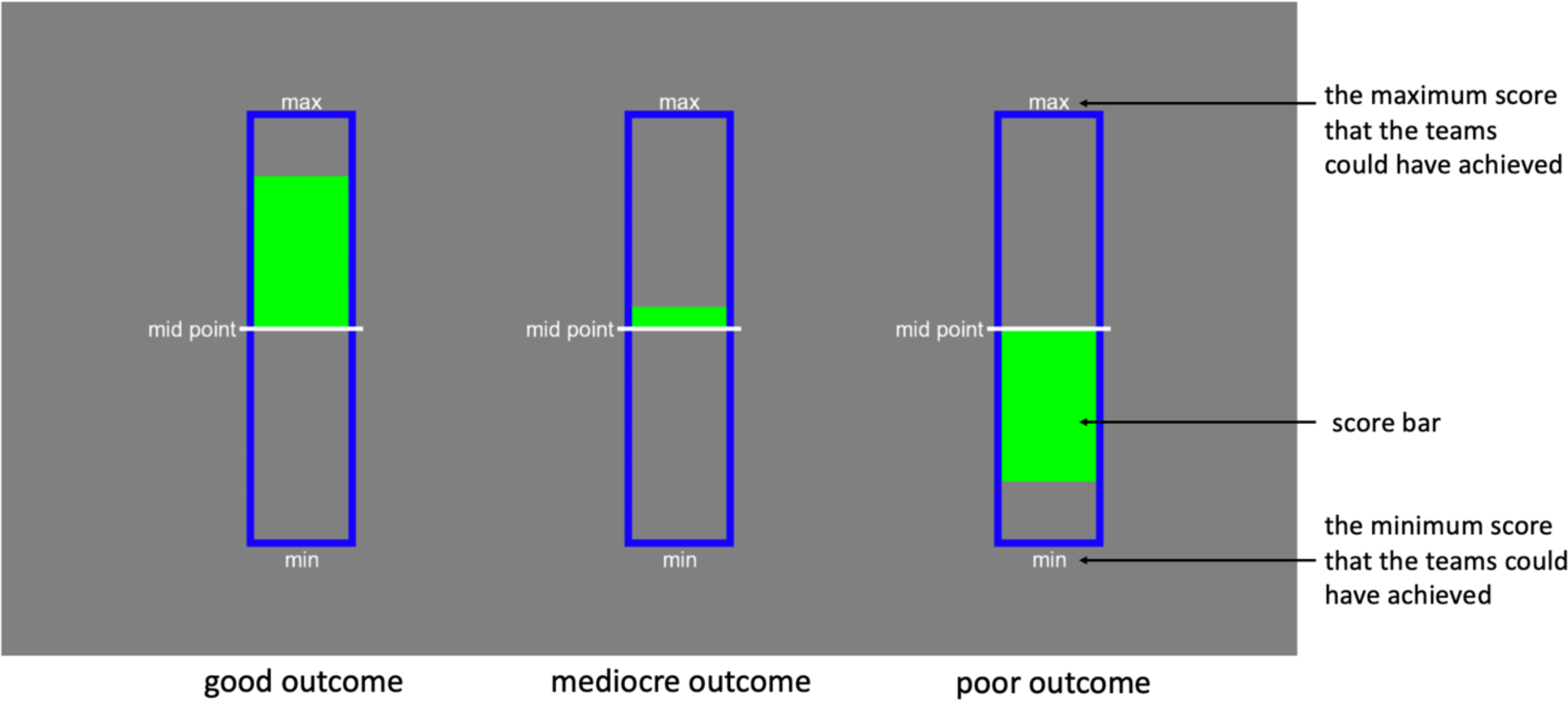

Then, you will be presented the photos of the two individuals in the team (Figure 3). One individual’s photo will be shown to you first either on the left (see example 1 in Figure 3) or right side (see example 2 in Figure 3) of the score frame and then after a few seconds the second individual’s photo will be shown on the other side. Then, you will be asked to decide which individual is more responsible for the outcome you just observed. You can choose the characters shown on the left or right sides by pressing the left and the right buttons, respectively. Your payment at the end of the experiment will depend on how well you predict the individual who was more responsible for a good or a bad outcome.

**Figure 6.**
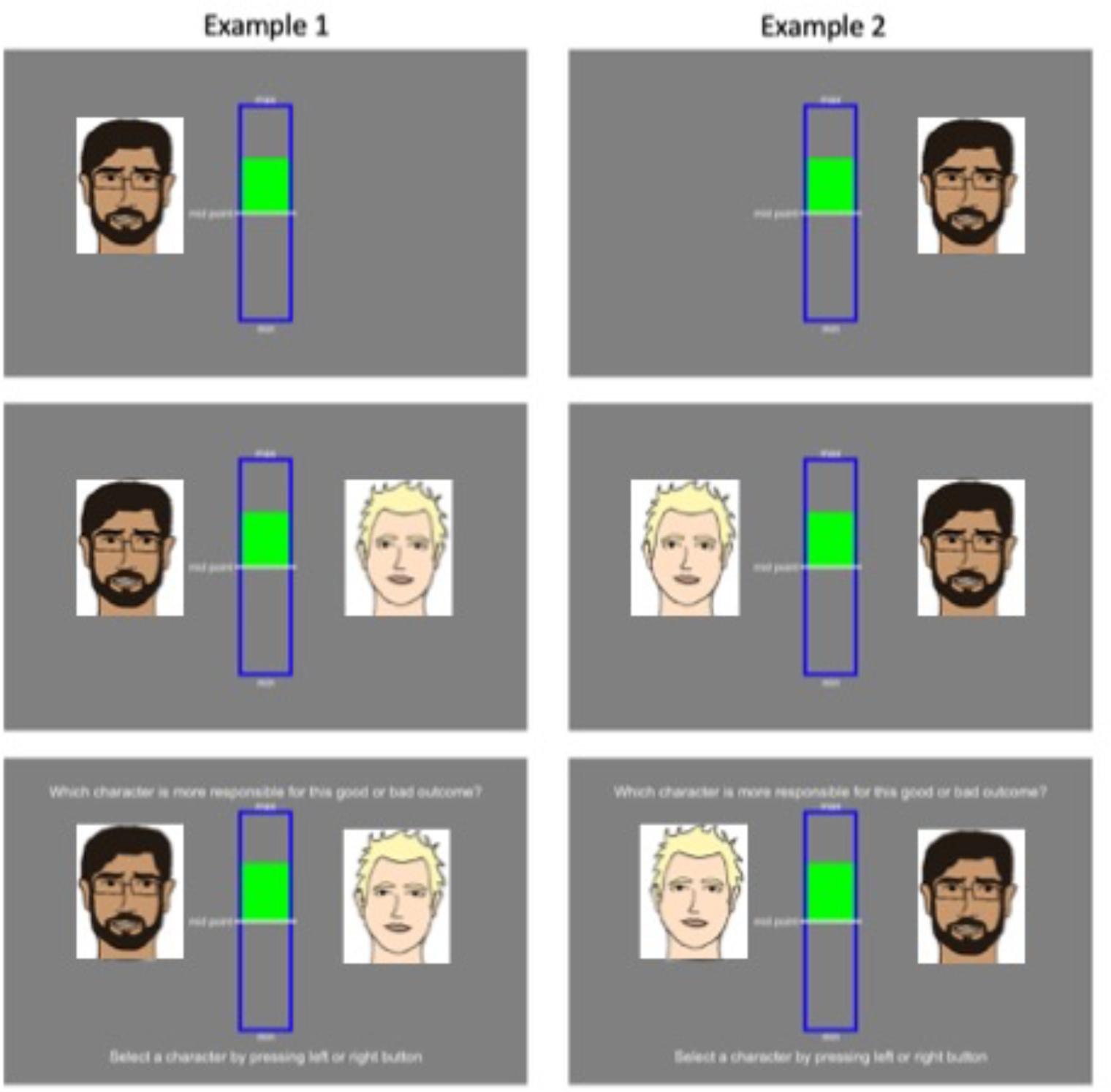

Once the task begins, you will be periodically asked to rate different individuals’ score/competence in the task. To do that, you will be shown an individual’s photo. Right below the photo, there is a tool to rate the presented individual’s score. As mentioned before, in each round, you see an outcome which is the result of two individuals’ teamwork. Individuals vary in their ability in the task. However, their performance might change from round to round due to natural fluctuation as they completed the minitask over several days. Therefore, it is important for you to track each individual’s score and do your rating based on that.

Your rating will be based on a scale ranging from min score to max score (seeing the rating scale and an example individual’s photo in Figure 4). Select your own answer by moving the scale using left and right buttons and lock your answer with the middle button on the response box. Moving the cursor all the way to the utmost left means that you think this individual is very poor at the task. However, moving the cursor all the way to the utmost right means that you think the individual is very competent at the task. The initial position of the cursor is always in the midpoint. You should move the cursor to where you think best describe the individual’s performance. Not moving the cursor will result in a severe loss of bonus payment.

**Figure 7.**
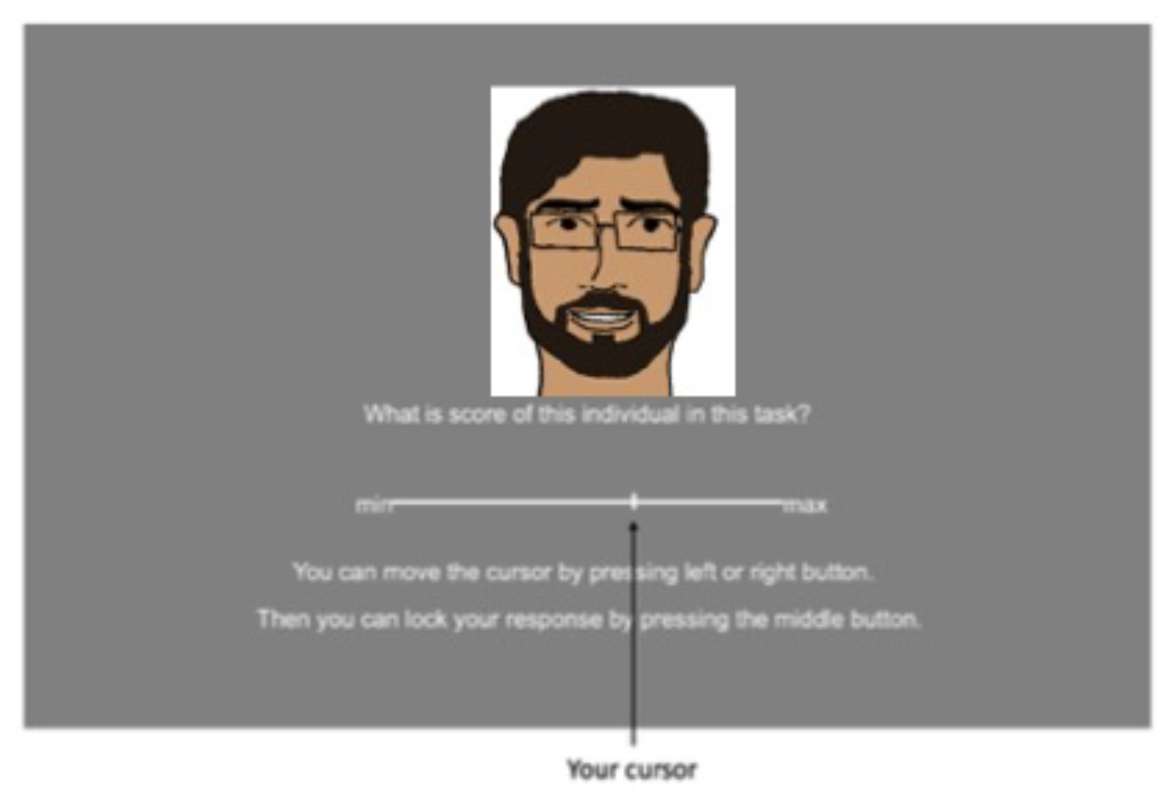

#### Experimental Procedure

You will complete the whole task in the fMRI scanner. The task will take around 60 minutes. We will also take a high-quality image of your brain during which you do not have to do anything but stay still in the scanner.

At the end of the task fMRI, you will be presented with images of different characters one after another. While at this stage you do not need to do anything, but you need to pay attention to the images that appear on the screen (Figure 5). Please keep your eyes on the fixation point during the interval between different images. At times, you will be asked to indicate the character which was just presented (Figure 6).

**Figure 8.**
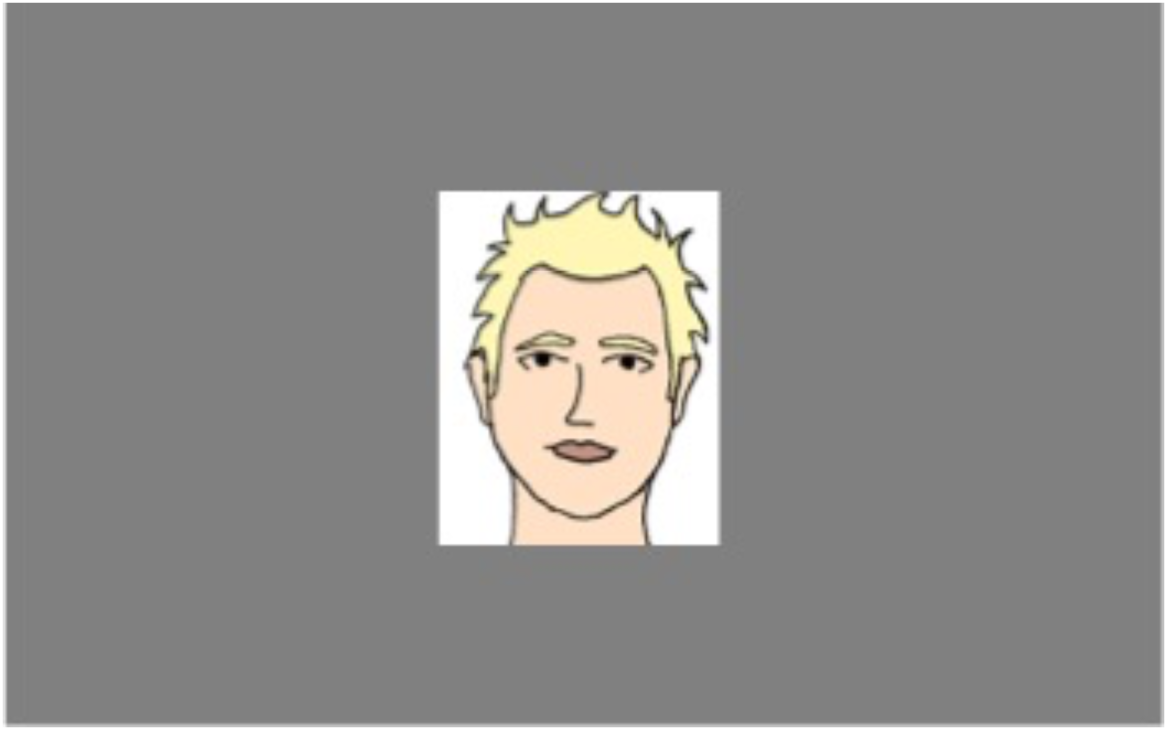

**Figure 9.**
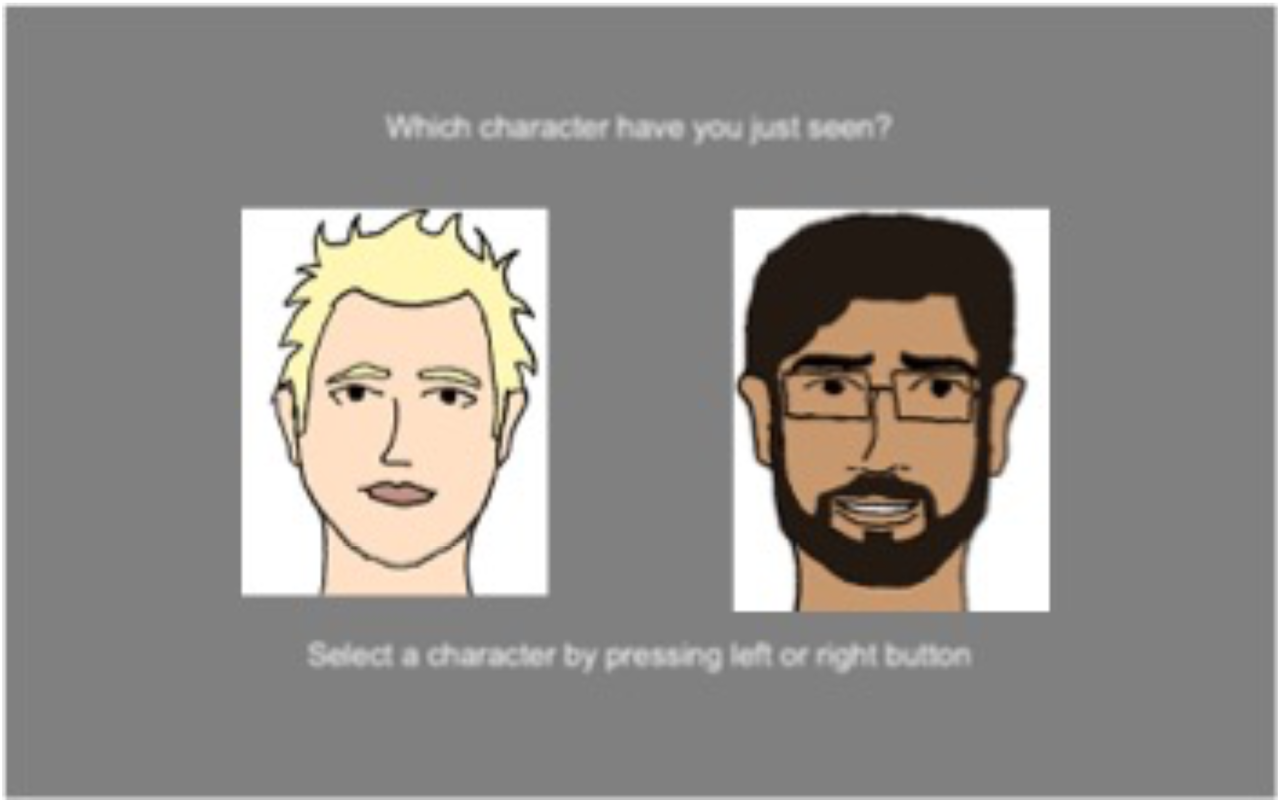

**Important note: in order to avoid gender bias in the social comparisons, we have chosen female and male characters for female and male participants, respectively.**

**Please let me know if you have any questions regarding the experiment.**

**--------------**

**END OF THE INSTRUCTION**

